# The CCL17-CCR4 axis is critical for mutant STAT6-mediated microenvironmental remodelling and therapeutic resistance in Relapsed/Refractory Diffuse Large B Cell Lymphoma

**DOI:** 10.1101/2024.12.13.628396

**Authors:** Madelyn J. Abraham, Cynthia Guilbert, Natascha Gagnon, Christophe Goncalves, Alexandre Benoit, Ryan N. Rys, Samuel E. J Preston, Ryan D. Morin, Wilson H. Miller, Nathalie A. Johnson, Sonia V. del Rincon, Koren K. Mann

## Abstract

Relapsed and refractory Diffuse Large B Cell Lymphoma (rrDLBCL) presents a significant challenge in hematology-oncology, with approximately 30-40% of DLBCL patients experiencing relapse or resistance to treatment. This underscores the urgent need to better understand the molecular mechanisms governing therapeutic resistance. Signal Transducer and Activator of Transcription 6 (STAT6) has been previously identified as a gene with recurrent D419 gain-of-function mutations in rrDLCBL. When STAT6^D419^ mutations are present in DLBCL tumour cells, we have demonstrated that transcription of the chemokine CCL17 (aka TARC) is increased, and tumours have increased infiltration of CD4+ T cells. However, the significance of increased T cell infiltration had not been determined. In the present study, we developed a mouse model of STAT6^D419N^ mutant DLBCL, that recapitulates the critical features of human STAT6^D419^ mutant DLBCL, including increased expression of phospho-STAT6, increased CD4+ T cell invasion, and resistance to doxorubicin treatment. With this model, we found CD4+ T cells in STAT6^D419N^ tumours have higher expression of the receptor for CCL17, CCR4. Using *ex vivo* functional assays we demonstrate that STAT6^D419N^ tumour cells are directly chemoattractive to CCR4+ CD4+ T cells, and when CCR4 is inhibited using a small molecule antagonist, CD4+ T cells in STAT6^D419N^ tumours are reduced and STAT6^D419N^ tumours regain therapeutic sensitivity to doxorubicin. Using PhenoCycler imaging of human rrDLBCL samples, we find that STAT6^D419^ tumours indeed have increased expression of phospho-STAT6+ and increased cellular interactions between phospho-STAT6+ tumour cells and CD4+/ CCR4+ CD4+ T cells. Thus, our data identify CCR4 as an attractive therapeutic target in STAT6^D419^ mutant rrDLBCL.

## Introduction

Diffuse Large B Cell Lymphoma (DLBCL) is the most common lymphoma in adults. Frontline treatment consists of poly-chemoimmunotherapy, most commonly rituximab, cyclophosphamide, doxorubicin, vincristine, and prednisone (R-CHOP), which leads to complete remission in 60% of patients. DLBCL that relapses or is refractory to R-CHOP (relapsed/refractory DLBCL: rrDLBCL) is associated with a poor outcome. Thus, it is critically important to better understand mechanisms of therapeutic resistance, with the overall goal of identifying treatment modalities with increased efficacy for rrDLBCL patients.

One interesting target in rrDLBCL is Signal Transducer and Activator of Transcription 6 (STAT6). *STAT6* is mutated or overactivated in several hematological malignancies, including cutaneous T cell lymphoma (CTCL) (1), primary mediastinal B cell lymphoma (PMBCL) (2–5), follicular lymphoma (FL) (6–9), and classical Hodgkin’s lymphoma (cHL) (10–12). In these cancer types, STAT6 signaling promotes tumour cell survival, proliferation, and resistance to apoptosis, leading to the classification of STAT6 as a potential therapeutic target in lymphoma. In DLBCL, STAT6 is not frequently overactivated, except in the case of relapsed/refractory disease, where our group has previously reported that mutations at the D419 gain-of-function hotspot are enriched (13). Indeed, we have shown that the presence of *STAT6^D419^* mutation in DLBCL cell lines leads to increased IL-4/13-dependent phospho-STAT6 nuclear residency. This correlates with increased expression of STAT6 transcriptional targets, including the chemokine *CCL17* (aka TARC), which is chemoattractive to CCR4+ immune cells, including CD4+ T cells. Concordantly, in human rrDLBCL, we found that increased tumour-cell derived *CCL17* is associated with increased CD4+ T cell invasion, specifically in STAT6-mutant samples (14).

Overall, research from our group and others has demonstrated that hyperactivation of STAT6, whether resulting from increased levels of IL-4/13 in the tumour microenvironment (TME) or due to gene mutation, can lead to tumour cell autonomous and microenvironmental changes. However, the clinical significance of these findings in rrDLBCL remains unclear. For instance, it is unknown if the CD4+ T cell infiltrate in *STAT6^D419^*-mutant tumours is skewed towards a specific polarization state or if these T cells contribute to disease progression and therapeutic resistance. Moreover, while we hypothesize that CCL17 is the main chemoattractant driving CD4+ T cell recruitment, this has not been experimentally proven. Recent publications have asserted that the spatial organization of tumours is just as important as their cellular constitution (15), but it is unknown if or how *STAT6^D419^* mutation induces tumour restructuring, and if CCL17-CCR4 plays a role. Finally, STAT6^D419^ mutations are enriched in rrDLBCL, but *in vitro* data indicates that STAT6^D419^ mutant tumour cells do not have increased resistance to any of the individual components of R-CHOP (14). It is still unknown if *STAT6^D419^*-mutations might contribute to therapeutic resistance due to changes in the TME.

In this study, we aimed to develop an immunocompetent mouse model that could recapitulate critical features of *STAT6^D419^*-mutant DLBCL, including increased phospho-STAT6 positivity in tumour tissue and increased CD4+ T cell invasion. With this model, we could address the outstanding questions related to *STAT6^D419^*-driven microenvironmental modification. Using flow cytometry and *ex vivo* functional assays, we demonstrate that *STAT6^D419N^*-mutant tumour cells are more chemoattractive to CCR4+ CD4+ T cells. Additionally, we show that STAT6^D419N^ tumours are indeed resistant to doxorubicin. Importantly, we demonstrate that blocking CCL17-induced CD4+ T cell recruitment to the STAT6^D419N^ TME, using a small molecule inhibitor of CCR4, re-sensitizes tumours to doxorubicin treatment. In human STAT6^D419^-mutant DLBCL biopsy tissues, we show that tumour cells have increased expression of phospho-STAT6 and CCL17. Moreover, STAT6^D419^-mutant tissues have increased interactions between phospho-STAT6+ tumour cells and CD4+/ CCR4+ CD4+ T cells as compared to STAT6^WT^ rrDLBCL samples. Thus, our data identifies CCR4 as an attractive therapeutic target for rrDLBCL patients with overactivated STAT6 signaling.

## Results

### The Immunocompetent mSTAT6^D419N^-Eµ-Myc Lymphoma Closely Recapitulates Critical Features of STAT6^D419^ Mutant rrDLBCL

Using *in vitro* models, we have previously demonstrated that STAT6^D419^ lymphoma cells have a similar proliferative rate and therapeutic sensitivity as STAT6^WT^ lymphoma cells (14). However, our previous data also indicated that *STAT6^D419^*mutation may function to induce therapeutic resistance via modulation of the TME. Thus, we developed an immunocompetent mouse model of STAT6^D419N^ lymphoma, to study how mutated *STAT6* impacts microenvironmental dynamics. To do this, we generated Eµ-Myc lymphoma cell lines overexpressing either murine (m) *STAT6^WT^* or *STAT6^D419N^*. These lymphoma cells can be injected into C57BL/6 mice via tail vein, where they home to the lymph nodes and spleen, and produce an orthotopic model of B cell lymphoma (**Figure 1A**). Disease burden can be followed via ultrasound (**Figure 1B**), and mice reach endpoint at 14-days post-injection, when cervical lymph node (cLN) size has exceeded 100 mm^3^. Critically, both Eµ-Myc-mSTAT6^WT^ tumours and Eµ-Myc-mSTAT6^D419N^ tumours grow at the same rate (**Figure 1C**), and neither genotype provides a survival advantage in the context of primary disease.

**Figure 1.**
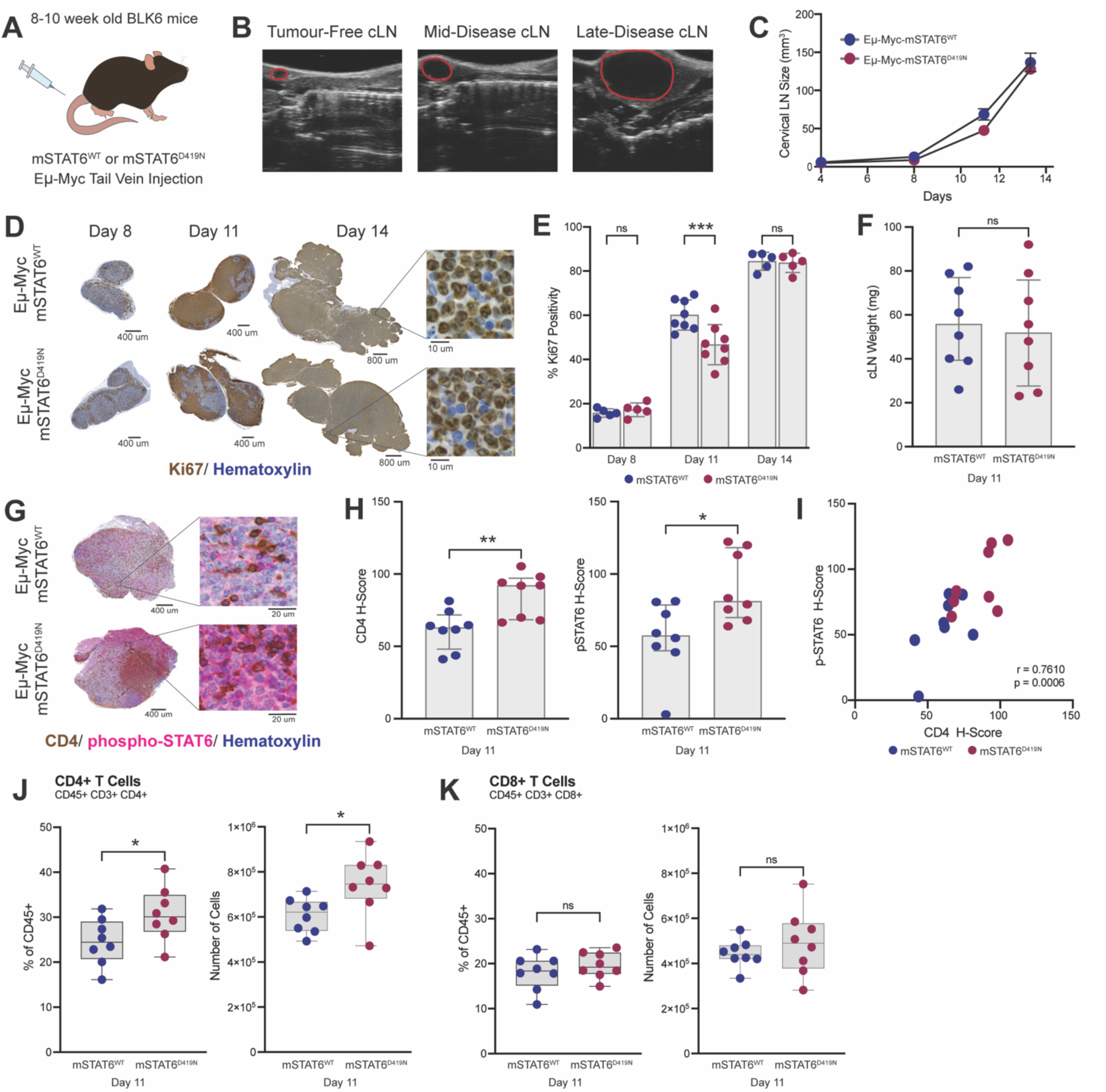
STAT6^D419N^-Eµ-Myc murine lymphoma tumours recapitulate critical features of human STAT6^D419^ lymphoma. **A.** To study the impact of STAT6^D419^ mutation on the lymphoma TME, 8-10 week old female C57BL/6 mice were injected via tail vein with mSTAT6^WT^ or mSTAT6^D419N^ Eµ-Myc tumour cells. **B.** Ultrasound images of tumour-free, mid-disease (day 11 post-injection), and late disease (day 14 post-injection) Eµ-Myc tumours, in the cLN. The anatomical structure that looks like a hatched line is the trachea. In all images, the cLN/tumour is outlined in red. **C.** cLN volume, quantified by ultrasound imaging, over time in mSTAT6^WT^ or mSTAT6^D419N^ Eµ-Myc tumour bearing mice. **D.** Representative Ki67 staining of early-, mid-, and late-disease mSTAT6^WT^ and mSTAT6^D419N^ Eµ-Myc tumours (day 8, day 11, and day 14, respectively). **E.** Quantification of Ki67 positive cells in mSTAT6^WT^ and mSTAT6^D419N^ Eµ-Myc tumours. At day 11, mSTAT6^D419N^ Eµ-Myc tumours have significantly fewer Ki67+ cells. **F.** Weights of mSTAT6^WT^ and mSTAT6^D419N^ Eµ-Myc tumours at day 11. Concordant with ultrasound data, mSTAT6^WT^ and mSTAT6^D419N^ cLN tumours are the same weight at this time point. **G.** Representative images of CD4 (brown) and phospho-STAT6 (pink) IHC co-staining in day 11 mSTAT6^WT^ and mSTAT6^D419N^ Eµ-Myc tumours. CD4 staining is consistently membranous, while phospho-STAT6 is expressed in both the cell cytoplasm and nucleus. **H.** Quantification of CD4 and phospho-STAT6 H-scores (see methods) in day 11 mSTAT6^WT^ and mSTAT6^D419N^ Eµ-Myc tumours. mSTAT6^D419N^ Eµ-Myc tumours have significantly increased CD4 and phospho-STAT6 expression. **I.** Correlation of CD4 and phospho-STAT6 expression in each tissue. Pearson correlation r = 0.7610. **J.** Quantification of CD4+ T cells in day 11 mSTAT6^WT^ and mSTAT6^D419N^ Eµ-Myc tumours. CD4+ T cells are live, single cells, that are CD45+, CD3+, and CD4+. Data is expressed as a percentage of CD45+ cells in each tissue, and as the total number of CD4+ T cells in each tissue. **K.** Quantification of CD8+ T cells in day 11 mSTAT6^WT^ and mSTAT6^D419N^ Eµ-Myc tumours. CD8+ T cells are live, single cells, that are CD45+, CD3+, and CD8+. Data is expressed as a percentage of CD45+ cells in each tissue, and as the total number of CD8+ T cells in each tissue.

To characterize these tumours over their development, lymph nodes were harvested at early-, mid-, and late-disease (days 8, 11, and 14 post-injection, respectively). At early and late disease, Eµ-Myc-mSTAT6^WT^ and Eµ-Myc-mSTAT6^D419N^ tumours had a similar percentage of Ki67+ cells within the lymph nodes (**Figure 1D-E**). However, at mid-disease, Eµ-Myc-mSTAT6^D419N^ tumours had significantly less Ki67+ tumour cells, despite no difference in tumour weight (**Figure 1F**). To characterize this discrepancy, we performed IHC co-staining for CD4 and phospho-STAT6 (**Figure 1G**). Consistent with patient samples, we found that Eµ-Myc-mSTAT6^D419N^ tumours had increased CD4 and phospho-STAT6 positivity (**Figure 1H**), and these two metrics were positively correlated (**Figure 1I**). This could be further recapitulated with flow cytometry (antibodies used for flow cytometry found in **Table S1**), where we found that Eµ-Myc-mSTAT6^D419N^ tumours were composed of an increased proportion of CD4+ T cells (represented as % of CD45+ cells within the tumour mass) and had a significantly increased total abundance of CD4+ T cells (**Figure 1J**). These differences were not observed at early or late disease (**Figure S1A-B**). No differences in proportion or abundance of CD8+ T cells were observed at any timepoint (**Figure 1K, Figure S1C-D**). Thus, our mouse model of STAT6^D419N^-mutant lymphoma closely replicates the human disease it is designed to model.

### mSTAT6^D419N^-Eµ-Myc Tumours have Increased CCR4+ Th1 Cells and Increased Hallmarks of Inflammation

We next questioned how CD4+ T cells are being enhanced in the Eµ-Myc-STAT6^D419N^ TME. It is well-established that the STAT6 transcriptional target CCL17 is overexpressed by *STAT6^D419^*-mutant lymphoma cells in response to IL-4 stimulation (6, 7, 14), and CCL17 can drive recruitment of CCR4+ immune cells, including CD4+ T cells. We first examined the expression of CCR4 on CD4+ T cells in Eµ-Myc-mSTAT6^WT^ and mSTAT6^D419N^ day 11 tumours. Flow cytometry analysis revealed a significant increase in the total number of CCR4+ CD4+ T cells in mSTAT6^D419N^ tumors compared to mSTAT6^WT^ tumors (**Figure 2A**). This suggests that mSTAT6^D419N^ tumors more effectively recruit or retain CCR4+ CD4+ T cells, consistent with the previously described increase in CCL17 in *STAT6^D419^* tumours (7, 14).

**Figure 2.**
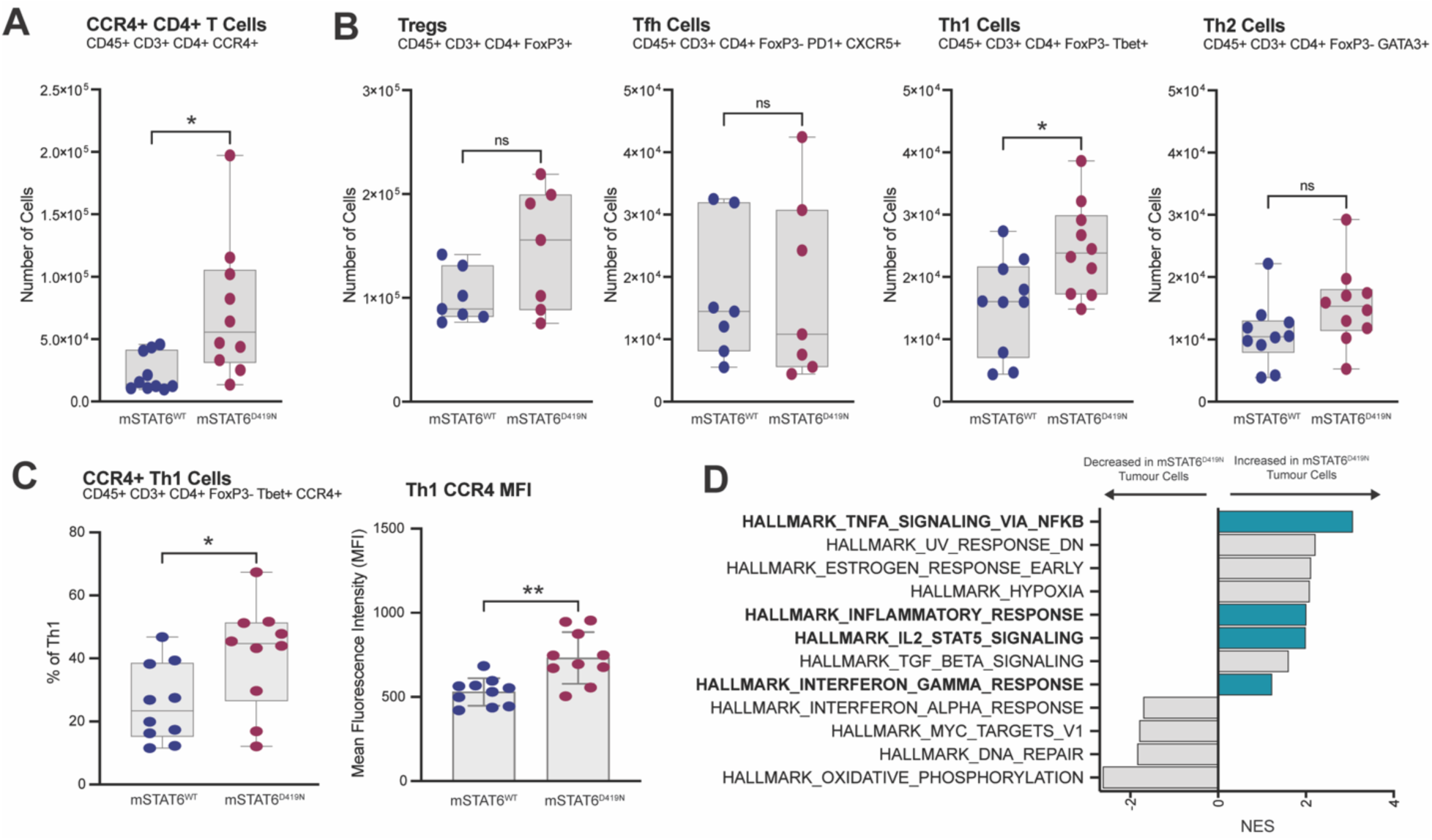
STAT6^D419N^-Eµ-Myc tumours have increased prevalence of CCR4+ Th1 cells. **A.** Quantification of CCR4+ CD4+ T cells in day 11 mSTAT6^WT^ and mSTAT6^D419N^ Eµ-Myc tumours. CCR4+ CD4+ T cells are live, single cells, that are CD45+, CD3+, CD4+, and CCR4+. Data is expressed as a total number of CCR4+ CD4+ T cells in each tissue. **B.** Quantification of Tregs (CD45+ CD3+ CD4+ FoxP3+), Tfh cells (CD45+ CD3+ CD4+ FoxP3-PD1+ CXCR5+), Th1 cells (CD45+ CD3+ CD4+ FoxP3-Tbet+), and Th2 (CD45+ CD3+ CD4+ FoxP3-GATA3+) in in day 11 mSTAT6^WT^ and mSTAT6^D419N^ Eµ-Myc tumours. Data is expressed as a total number of cells in each tissue. **C.** Expression of CCR4 on Th1 cells in day 11 mSTAT6^WT^ and mSTAT6^D419N^ Eµ-Myc tumours, expressed as total percentage of Th1 cells that are CCR4+ (CD45+ CD3+ CD4+ FoxP3-Tbet+ CCR4+), and as the mean fluorescence intensity (MFI) of CCR4 expression on Th1 cells. **D.** Gene set enrichment analysis (GSEA) obtained from RNAseq of isolated tumour cells (ie. CD19+ cells) from day 11 mSTAT6^WT^ and mSTAT6^D419N^ Eµ-Myc tumours. Teal colouring indicates enrichment of pathways associated with inflammation and inflammatory signaling. NES = Negative Enrichment Score.

We further compared the polarization states of CD4+ T cells in mSTAT6^WT^ and mSTAT6^D419N^ tumours by quantifying regulatory T cells (Tregs), T follicular helper cells (Tfh), inflammatory T helper cells (Th1), and anti-inflammatory T helper cells (Th2) using flow cytometry (**Figure 2B**). Among these subsets, only Th1 cells were significantly increased in mSTAT6^D419N^ tumors, indicating a skewed immune profile favoring Th1 cell recruitment or expansion. Indeed, a higher percentage of Th1 cells in mSTAT6^D419N^ tumors were CCR4-positive compared to those in mSTAT6^WT^ tumors. Additionally, Th1 cells in mSTAT6^D419N^ tumors exhibited a significantly higher mean fluorescence intensity (MFI) of CCR4 expression (**Figure 2C**).

To explore the broader impact of immune changes on the tumour compartment, we isolated Eµ-Myc cells from mSTAT6^WT^ and mSTAT6^D419N^ mid-disease tumours, using CD19 microbeads, and performed bulk RNA sequencing. Gene set enrichment analysis (GSEA) indicated that mSTAT6^D419N^ tumour cells had increased hallmarks of inflammation (**Figure 2D**), consistent with the elevated presence of Th1 cells. Of note, Th1 cells in mSTAT6^WT^ and mSTAT6^D419N^ tumours did not express more interferon-gamma (IFNγ), nor did they show increased expression of activation or degranulation markers (CD69 and CD107a, respectively; **Figure S2A-C**). Additionally, the expression of these markers was not changed on CD8+ T cells (**Figure S2D-F**). These findings demonstrate that mSTAT6^D419N^ mutation leads to the increased presence of CCR4+Th1 cells, that may contribute to an enhanced inflammatory response signature in mSTAT6^D419N^ tumour cells.

### mSTAT6^D419N^ Tumours are Chemoattractive to CCR4+ CD4+ T Cells

To experimentally validate that STAT6^D419N^ tumor cells attract CCR4+ immune cells, we conducted a series of *ex vivo* assays designed to assess the chemoattractive properties of Eµ-Myc-mSTAT6^WT^ and mSTAT6^D419N^ tumor cells. We first tested the ability of Eµ-Myc tumour cells in culture to attract various CCR4+ immune cells, including CD4+ T cells, eosinophils, dendritic cells, macrophages, NK cells, and CD8+ T cells. In this experiment, mSTAT6^WT^ and mSTAT6^D419N^ Eµ-Myc cells were plated in a 6-well dish and were stimulated with IL-4. One hour following stimulation, bulk splenocytes from non-tumour bearing mice were plated overtop the Eµ-Myc tumour cells, using a 5 µm pore membrane. Splenocytes were allowed to migrate towards the tumour cells for 16 hours, then migrated cells were collected and characterized by flow cytometry (**Figure 3A**). Concordant with our previous studies, qPCR confirmed that mSTAT6^D419N^ tumour cells produced significantly more *CCL17* in response to IL-4 stimulation than mSTAT6^WT^ tumour cells (**Figure 3B**). Indeed, IL-4-stimulated tumour cells induced significant migration of CCR4+ splenocytes, predominantly consisting of CD4+ T cells (**Figure 3C**). As predicted, STAT6^D419N^ cells induced significantly greater migration of CCR4+ CD4+ T cells as compared to STAT6^WT^ cells. This migration was blocked by pre-treating splenocytes with the CCR4 inhibitor AZD2098 (16). These findings demonstrate that STAT6^D419N^ tumor cells are more effective at attracting CCR4+ CD4+ T cells, highlighting the functional impact of increased CCL17 production by mSTAT6^D419N^ tumour cells.

**Figure 3.**
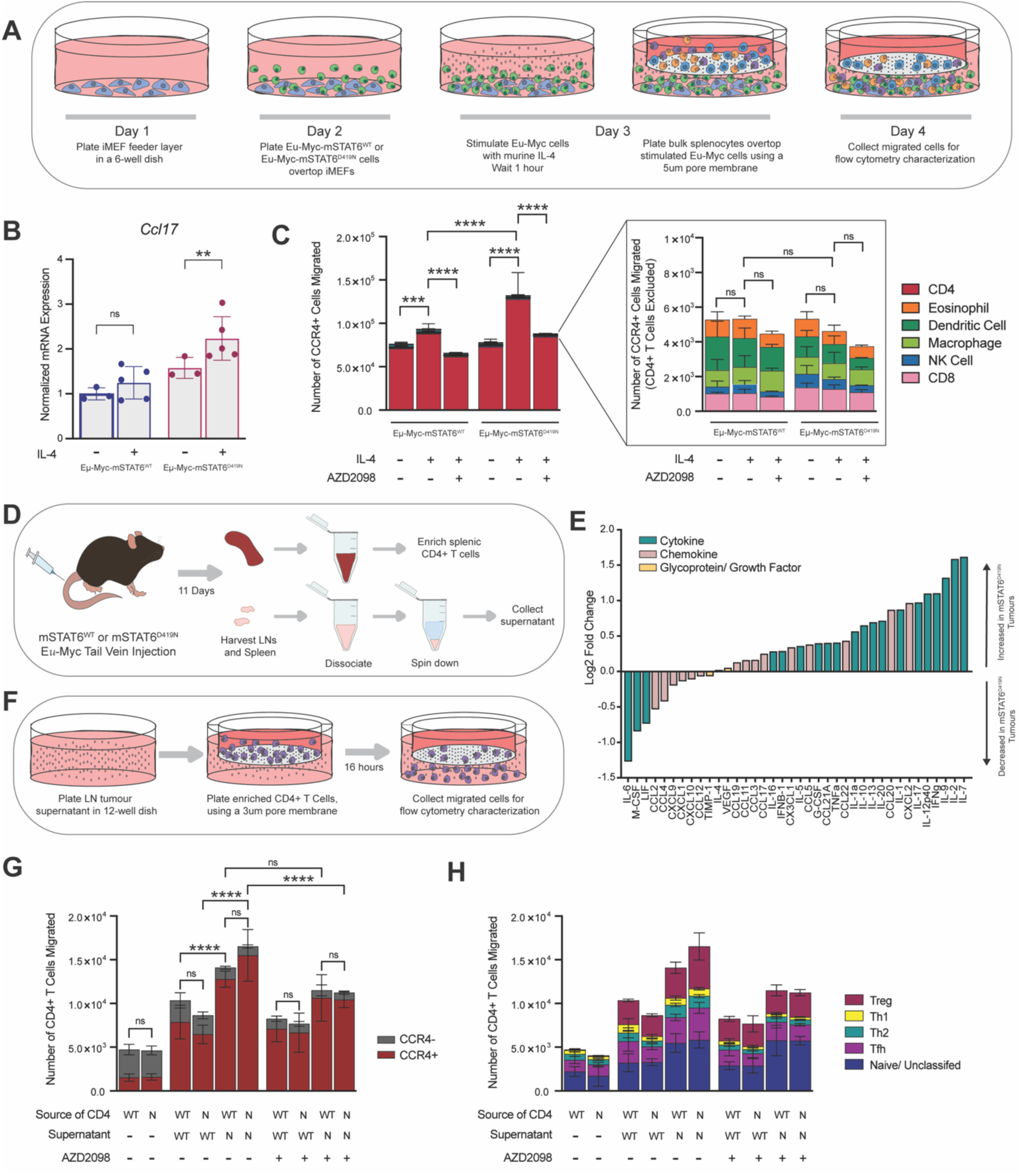
STAT6^D419N^-Eµ-Myc tumours are directly chemoattractive to CCR4+ CD4+ T cells. **A.** Schematic demonstrating the workflow for determining the ability of mSTAT6^WT^ and mSTAT6^D419N^ Eµ-Myc tumour cells, +/- mIL-4 to attract different cell types from tumour-naïve spleen, +/- AZD2098. **B.** Quantification of *Ccl17* expression in mSTAT6^WT^ and mSTAT6^D419N^ Eµ-Myc tumour cells, following 1 hour of mIL-4 or vehicle stimulation. **C.** Quantification of CCR4+ splenocytes that migrated towards mSTAT6^WT^ or mSTAT6^D419N^ Eµ-Myc tumour cells, as determined by flow cytometry. (**C_i_)** shows all migrated cells. (**C_ii_)** is the same data but excluding CD4+ T cells for better visualization of other migrated cellular subsets. **D.** Schematic demonstrating the workflow for collecting tumour-exposed CD4+ T cells from spleen and day 11 mSTAT6^WT^ and mSTAT6^D419N^ Eµ-Myc day 11 tumour supernatants. **E.** Waterfall plot showing the log2 fold change in expression of cytokines and chemokines between mSTAT6^WT^ and mSTAT6^D419N^ Eµ-Myc day 11 tumour supernatants. **F.** Schematic demonstrating the workflow for determining the ability of mSTAT6^WT^ and mSTAT6^D419N^ Eµ-Myc day 11 tumour supernatants to attract CD4+ T cells from tumour-exposed spleen. **G.** Quantification of CCR4+ CD4+ T cells enriched from either STAT6^WT^ or STAT6^D419N^ Eµ-Myc day 11 tumour bearing mice that migrated towards STAT6^WT^ or STAT6^D419N^ Eµ-Myc day 11 tumour supernatants. **H.** Data from (**G**), expressed as CD4+ T cell polarization state (ie. Treg, Th1, Th2, Tfh, or Naïve/ Unclassified).

*In vivo*, there is extensive crosstalk between tumour cells, immune cells, and stroma, that is not recapitulated with splenocytes from tumour-naïve mice migrating towards tumour cells stimulated with a single cytokine. Thus, we next attempted to more closely model the complexity of an intact TME *ex vivo*, to better characterize CD4+ T cell migration. To do this, mice were injected with either mSTAT6^WT^ or mSTAT6^D419N^ tumor cells, and at mid-disease, tumors and spleens were harvested and dissociated. CD4+ T cells were enriched from the tumour-bearing spleens, and tumor supernatants were collected from lymph nodes (**Figure 3D**). Thus, with this model, CD4+ T cells have been polarized due to tumour exposure, and tumour supernatants represent the secretome of an intact TME. Chemokine/cytokine profiling of tumour supernatants demonstrated that mSTAT6^D419N^ tumours tended to express higher levels of many cytokines/chemokines, including Th1-associated cytokines IFNγ, IL-2, TNFα, and IL-12, and the CCR4-ligands and STAT6 transcriptional targets CCL17 and CCL22 (**Figure 3E**). Supernatants from either mSTAT6^WT^ or mSTAT6^D419N^ tumours were then used as chemoattractants to CD4+ T cells derived from tumour-bearing spleens (**Figure 3F**).

We found that significantly more CCR4+ CD4+ T cells migrated towards STAT6^D419N^ supernatant compared to STAT6^WT^ supernatant, regardless of whether the CD4+ T cells were derived from STAT6^WT^ or STAT6^D419N^ tumour-bearing mice (**Figure 3G**). This migration was markedly reduced when CD4+ T cells were pretreated with AZD2098, indicating the role of CCR4-CCL17 interaction in transmigration of CD4+ T cells towards tumour supernatant. We further phenotyped the migrated T cells, and found that similar proportions of Treg, Tfh, Th1, Th2, and naive/unclassified CD4+ T cells migrated towards both STAT6^WT^ and STAT6^D419N^ supernatants (**Figure 3H**), suggesting that the chemoattractive effect of STAT6^D419N^ supernatant is not specific to any CD4+ T cell subset. These experiments collectively demonstrate that STAT6^D419N^ tumors possess a heightened ability to attract CCR4+ CD4+ T cells through increased production of CCL17, thereby enhancing the CD4-rich microenvironmental characteristics of STAT6^D419N^ tumours.

### mSTAT6^D419N^ Eµ-Myc Tumours are Resistant to Doxorubicin Treatment

*In vitro,* mSTAT6^D419N^ Eµ-Myc tumour cells do not have altered sensitivity to any of the individual components of R-CHOP (17). However, our data indicate that *mSTAT6^D419N^* mutation drives increased frequency of CCR4+ CD4+ T cells, leading to evidence of enhanced tumour cell inflammatory signaling, which is known to correlate with doxorubicin resistance in human DLBCL patients (18–21). Thus, we hypothesized that an intact tumour microenvironment is required to truly realize the impact of *STAT6^D419^* mutation on therapeutic response, and that our mouse model of *STAT6^D419^*-mutant lymphoma may not recapitulate the results previously obtained *in vitro*.

To investigate this, mice were injected with Eµ-Myc mSTAT6^WT^ or mSTAT6^D419N^ tumour cells on day 1, then were given 3 mg/kg of doxorubicin at day 11, day 13, and day 15 post-injection (**Figure 4A**). Over the course of disease, tumours were monitored via ultrasound. We found that mSTAT6^WT^ tumours returned to baseline size by day 15 post-injection, indicating the efficacy of doxorubicin treatment. In contrast, mSTAT6^D419N^ tumour volume stabilized during doxorubicin treatment, but tumours continued to grow immediately upon the cessation of treatment (**Figure 4B**). Additionally, ultrasound monitoring of the secondary tumour site, the inguinal lymph node (iLN), revealed that 100% of mice with mSTAT6^WT^ tumours had sustained disease clearance in the iLN, whereas 40% of mice with mSTAT6^D419N^ tumours had bilateral iLN relapse (**Figure 4C**). While not achieving statistical significance (Gehan-Breslow-Wilcoxon test p = 0.0764), this led to decreased overall survival of mSTAT6^D419N^ tumour bearing mice treated with doxorubicin (**Figure 4D**).

**Figure 4.**
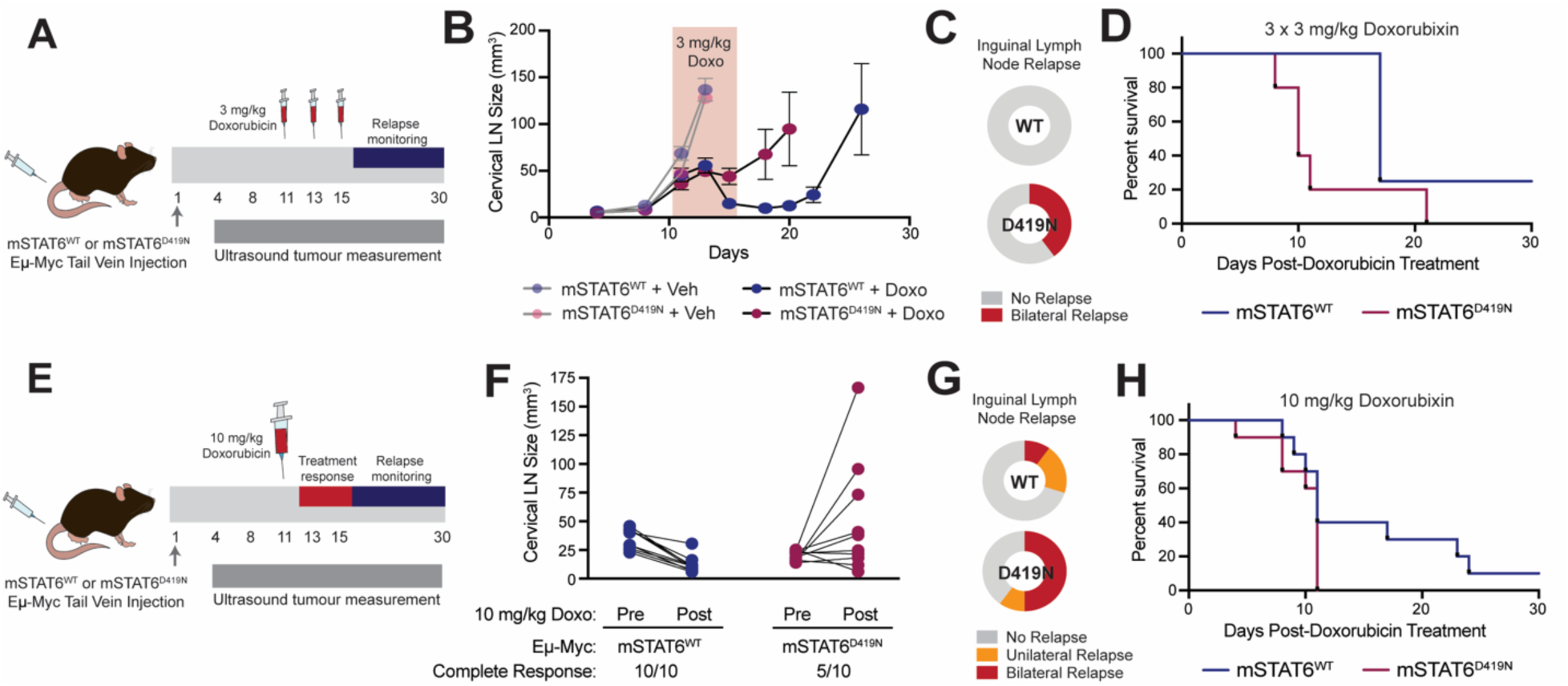
mSTAT6^D419N^-Eµ-Myc tumours are resistant to doxorubicin treatment. **A.** Schematic demonstrating the timeline for 3 x 3 mg/kg doxorubicin treatment and relapse monitoring for mSTAT6^WT^ and mSTAT6^D419N^ Eµ-Myc tumour bearing mice. **B.** Growth curve of mSTAT6^WT^ and mSTAT6^D419N^ Eµ-Myc tumours, treated with either 3 x 3 mg/kg doxorubicin or vehicle control. Red shaded area indicates the days where doxorubicin treatment was given. **C.** Pie charts showing the proportion of mice with secondary tumour site (ie. the iLN) relapse following 3 x 3 mg/kg doxorubicin treatment. **D.** Kaplan-Meier survival curve of mSTAT6^WT^ and mSTAT6^D419N^ Eµ-Myc tumour bearing mice following 3 x 3 mg/kg doxorubicin treatment. **E.** Schematic demonstrating the timeline for 10 mg/kg doxorubicin treatment and relapse monitoring for mSTAT6^WT^ and mSTAT6^D419N^ Eµ-Myc tumour bearing mice. **F.** Treatment response of mSTAT6^WT^ and mSTAT6^D419N^ Eµ-Myc tumour bearing mice following 10 mg/kg doxorubicin. Connecting lines show the pre- and post-treatment cLN volumes in individual mice. If the slope of the connecting line is negative, mice are considered to have had a complete response to doxorubicin. Pre-treatment is day 11 post-Eµ-Myc injection, and post-treatment is day 15 post-Eµ-Myc injection. **G.** Pie charts showing the proportion of mice with iLN relapse following 10 mg/kg doxorubicin treatment. **H.** Kaplan-Meier survival curve of mSTAT6^WT^ and mSTAT6^D419N^ Eµ-Myc tumour bearing mice following 10 mg/kg doxorubicin treatment.

We next changed our treatment paradigm, to more closely model primary-refractory disease, by giving each mouse a single high-dose doxorubicin treatment of 10 mg/kg at day 11 post-injection, which represents a timepoint where tumours have established, and our data indicates that mSTAT6^D419N^ tumours have CCR4+ CD4+ T cell invasion **(Figure 4E**). Similar to our findings with 3 x 3 mg/kg doxorubicin, we found that 100% of mice bearing mSTAT6^WT^ tumours decreased to baseline size, while only 50% of mice bearing mSTAT6^D419N^ tumours decreased in size (**Figure 4F**). This was recapitulated in the iLN, where 100% of mice bearing mSTAT6^WT^ tumours had durable disease clearance in the iLN, while 50% of mice bearing mSTAT6^D419N^ tumours had bilateral iLN relapse, and another 10% had unilateral iLN relapse (**Figure 4G**). Accordingly, mSTAT6^WT^ tumour bearing mice had a trend towards improved overall survival (Gehan-Breslow-Wilcoxon test p = 0.1211; **Figure 4H**).

Tumours collected at disease endpoint showed no difference in tumour burden or skewing in the proportions of CD4+ T cells, and CD8+ T cells between mSTAT6^WT^ and mSTAT6^D419N^ tumours (**Figure S3A-D**). These results indicate that the mSTAT6^D419N^-driven changes in the TME at day 11 are sufficient to engender resistance to doxorubicin, but that changes in the TME are not retained or exaggerated at disease relapse in this mouse model.

### Treatment of mSTAT6^D419N^ Eµ-Myc Tumours with the CCR4 Inhibitor AZD2098 Re-Sensitizes them to Doxorubicin

We next aimed to investigate how inhibiting CCR4 would affect the growth of Eµ-Myc tumors, following our discovery that *mSTAT6^D419N^* mutation in Eu-Myc tumour cells enhances the recruitment of CCR4+ CD4+ T cells. To do this, we treated Eµ-Myc tumor-bearing mice with AZD2098 and monitored tumor growth and changes in the TME over time (**Figure 5A**). AZD2098 was administered via oral gavage twice weekly, and we found that it successfully blocked the mid-disease invasion of CD4+ T cells into mSTAT6^D419N^ tumors (**Figure 5B**). Further analysis showed that AZD2098 did not affect the numbers of Tregs or Tfh cells in either mSTAT6^WT^ or mSTAT6^D419N^ tumors. However, as previously noted, Th1 cells were more abundant in mSTAT6^D419N^ tumors compared to mSTAT6^WT^, and treatment with AZD2098 resulted in a modest decrease in Th1 cells in mSTAT6^D419N^ tumors. Consistent with CCR4’s role in Th2 cells (22, 23), AZD2098 treatment reduced Th2 cells in both mSTAT6^WT^ and mSTAT6^D419N^ tumors (**Figure 5C**). Despite these shifts in TME composition, AZD2098 treatment did not significantly alter the growth rate of either mSTAT6^WT^ or mSTAT6^D419N^ tumors (**Figure 5D**).

**Figure 5.**
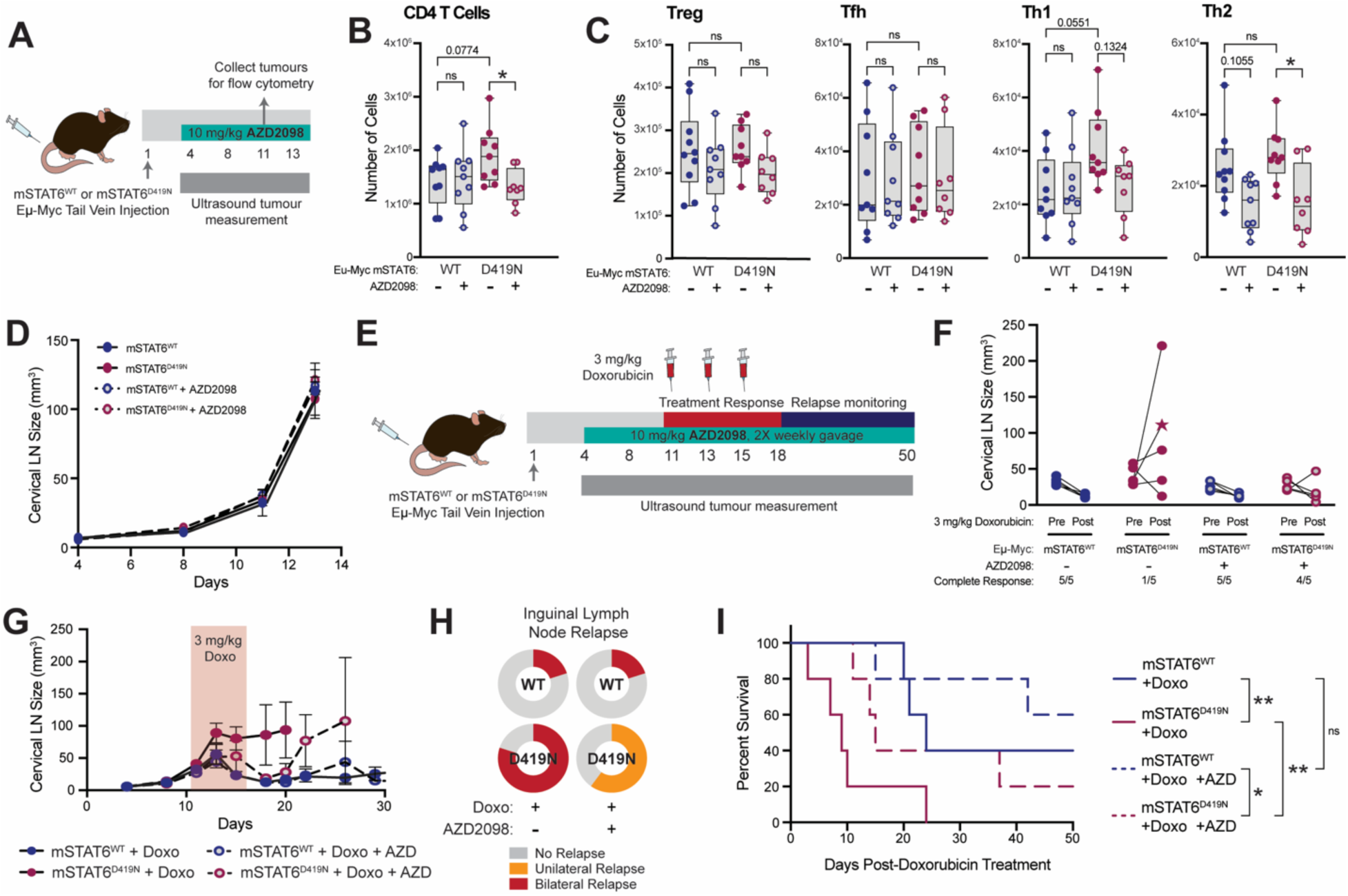
CCR4 inhibition resensitizes mSTAT6^D419N^-Eµ-Myc tumours to doxorubicin treatment. **A.** Schematic demonstrating the timeline for determining the impact of AZD2098 treatment on mSTAT6^WT^ and mSTAT6^D419N^ Eµ-Myc TME composition and tumour growth. **B.** Quantification of CD4+ T cells in mSTAT6^WT^ and mSTAT6^D419N^ day 11 Eµ-Myc tumours, following treatment with AZD2098 or vehicle control. **C.** Quantification of Tregs (CD45+ CD3+ CD4+ FoxP3+), Tfh cells (CD45+ CD3+ CD4+ FoxP3-PD1+ CXCR5+), Th1 cells (CD45+ CD3+ CD4+ FoxP3-Tbet+), and Th2 (CD45+ CD3+ CD4+ FoxP3-GATA3+) in mSTAT6^WT^ and mSTAT6^D419N^ day 11 Eµ-Myc tumours, following treatment with AZD2098 or vehicle control. **D.** Growth curve of mSTAT6^WT^ and mSTAT6^D419N^ Eµ-Myc tumours, treated with either 2 times weekly 10 mg/kg AZD2098 or vehicle control. **E.** Schematic demonstrating the timeline for 3 x 3 mg/kg doxorubicin and 10 mg/kg AZD treatment and relapse monitoring for mSTAT6^WT^ and mSTAT6^D419N^ Eµ-Myc tumour bearing mice. **F.** Treatment response of mSTAT6^WT^ and mSTAT6^D419N^ Eµ-Myc tumour bearing mice following 3 x 3 mg/kg doxorubicin and 10 mg/kg AZD2098. Connecting lines show the pre- and post-treatment cLN volumes in individual mice. Pre-treatment is day 11 post-Eµ-Myc injection, and post-treatment is day 18 post-Eµ-Myc injection. Star indicates the cLN volume is from day 15 post-Eµ-Myc injection, and the mouse did not survive until day 18 post-Eµ-Myc injection. **G.** Growth curve of mSTAT6^WT^ and mSTAT6^D419N^ Eµ-Myc tumours, treated with 3 x 3 mg/kg doxorubicin and 10 mg/kg AZD2098, or vehicle controls. Red shaded area indicates the days where doxorubicin treatment was given**. H.** Pie charts showing the proportion of mice with secondary tumour site (ie. the iLN) relapse following 3 x 3 mg/kg doxorubicin and 10 mg/kg AZD2098 treatment. **I.** Kaplan-Meier survival curve of mSTAT6^WT^ and mSTAT6^D419N^ Eµ-Myc tumour bearing mice following combination treatment with 3 x 3 mg/kg doxorubicin with 10 mg/kg AZD2098.

Encouraged by AZD2098’s ability to reduce CD4+ T cell infiltration in mSTAT6^D419N^ tumors to levels comparable to mSTAT6^WT^, we hypothesized that the addition of AZD2098 could restore doxorubicin sensitivity in mSTAT6^D419N^ tumors (**Figure 5E**). As in previous experiments, 100% of mSTAT6^WT^ tumors responded completely to doxorubicin alone, whereas only 20% of mSTAT6^D419N^ tumors responded (**Figure 5F**). Remarkably, combining AZD2098 with doxorubicin resulted in a complete response of 100% of mSTAT6^WT^ tumors and 80% of mSTAT6^D419N^ tumors. While mSTAT6^D419N^ tumors treated with doxorubicin alone resumed growth immediately after treatment cessation, tumors treated with the combination of doxorubicin and AZD2098 did not regrow until approximately one-week post-treatment (**Figure 5G**). Additionally, 80% of mSTAT6^D419N^ tumors treated with doxorubicin alone showed bilateral inguinal lymph node relapse, whereas those treated with the combination showed only unilateral relapse, indicating a minor reduction in systemic disease (**Figure 5H**). Overall, the combination treatment significantly improved survival in mice bearing mSTAT6^D419N^ tumors, but not in those with mSTAT6^WT^ tumors (**Figure 5I**). These findings demonstrate that inhibiting the CCL17-CCR4 signaling axis re-sensitizes mSTAT6^D419N^ Eµ-Myc tumors to doxorubicin treatment.

### PhenoCycler Imaging of STAT6^D419^-Mutant rrDLBCL Patient Samples Shows Increased Expression of phospho-STAT6, CCL17, and CCR4

To validate our results in human samples, we performed highly multiplexed PhenoCycler immunofluorescent imaging (24–26), on a DLBCL TMA with biopsy tissues collected at disease relapse (**Table S2**). We additionally included slides with full biopsy specimens from 6 patients with STAT6^D419^-mutant DLBCL, to compare how the presence of STAT6 mutation impacts the CCL17-CCR4 signaling axis (**Figure 6A-B**). Of these STAT6^D419^-mutant tissue biopsies, three were relapse biopsies from DLBCL patients, one was a diagnostic DLBCL biopsy from a patient who proceeded to relapse, one was a diagnostic primary mediastinal B cell lymphoma (PMBCL) biopsy, and one was a diagnostic follicular lymphoma biopsy, from a tumour that proceeded to transform into DLBCL (GCB) involving the central nervous system (DLBCL-CNS).

**Figure 6.**
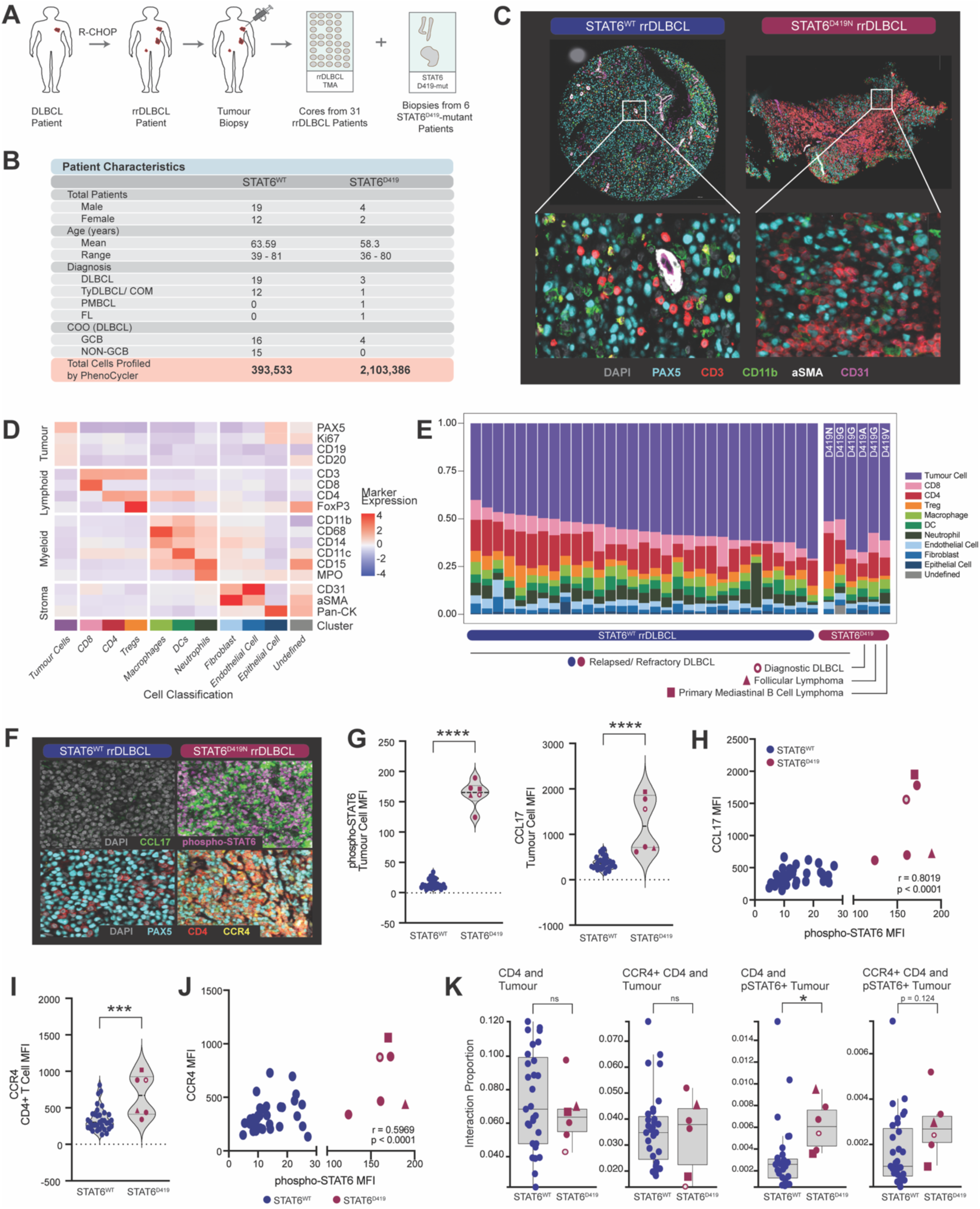
PhenoCycler Imaging of human rrDLBCL biopsies reveals increased phospho-STAT6, CCL17, and CCR4 in STAT6^D419^-mutant patients. **A.** Schematic depicting the human DLBCL samples used in this study for PhenoCycler imaging. A TMA was constructed with cores from rrDLBCL tissues. Additionally, whole biopsies from STAT6^D419^-mutant patients were imaged. **B.** Patient characteristics for the biopsies included in this study. **C.** Representative images of a STAT6^WT^ rrDLBCL TMA core, and a STAT6^D419N^ rrDLBCL biopsy. **D.** Heatmap showing the mean expression of different markers with cell classifications. **E.** Proportions of different cell types in STAT6^WT^ rrDLBCL and STAT6^D419^-mutant lymphoma biopsies. STAT6^D419^ biopsies included a diagnostic biopsy (open circle), a follicular lymphoma biopsy (triangle), and a primary mediastinal B cell lymphoma (square). **F.** Representative images of pSTAT6, CCL17, and CCR4 in a STAT6^WT^ and STAT6^D419N^ rrDLBCL biopsy. **G.** Mean fluorescence intensity (MFI) of phospho-STAT6 and CCL17 in tumour cells in STAT6^WT^ and STAT6^D419^-mutant biopsies. **H.** Correlation of phospho-STAT6 and CCL17 in each biopsy. Pearson correlation r = 0.8019. **I.** MFI of CCR4 in CD4+ T cells in STAT6^WT^ and STAT6^D419^-mutant biopsies. **J.** Correlation of phospho-STAT6 and CCR4 in each biopsy. Pearson correlation r = 0.5969. **K.** Interaction proportion between CD4+ T cells and tumour cells, CCR4+ CD4+ T cells and tumour cells, CD4+ T cells and phospho-STAT6+ tumour cells, and CCR4+ CD4+ T cells and phospho-STAT6+ T cells.

In our multiplexed images, we were clearly able to visualize tumour cells, T cells, myeloid cells, fibroblasts, and endothelial cells within human DLBCL tissues (**Figure 6C, Figure S4**). Tumour tissues were analyzed by segmenting images into cells, then performing unsupervised clustering based on cellular expression of the lineage markers CD11b, CD11c, CD14, CD15, CD20, CD21, CD31, CD3, CD4, CD8, CD68, FoxP3, Ki67, MPO, PAX5, pan-CK, and αSMA (**Figure 6D**). We then compared the proportion of these different cell types in rrDLBCL and STAT6^D419^-mutant samples (**Figure 6E**). Concordant with our previous observations (14), STAT6^D419^-mutant rrDLBCL samples were among the highest expressors of CD4+ T cells. Intriguingly, the STAT6^D419^-mutant samples from diagnostic DLBCL, FL, and PMBCL had relatively fewer CD4+ T cells.

We next examined phospho-STAT6 and CCL17 expression in tumour cells, and CCR4 expression in CD4+ T cells (**Figure 6F**). In tumour cells in STAT6^D419^-mutant samples, phospho-STAT6 and CCL17 expression were significantly increased, as measured by mean fluorescence intensity (MFI) of each marker (**Figure 6G**). Additionally, the expression of CCL17 and phospho-STAT6 were significantly positively correlated (**Figure 6H**). CCR4 showed a similar increase in expression in CD4+ T cells from STAT6^D419^-mutant patients, and this was also significantly positively correlated with phospho-STAT6 expression (**Figure 6I-J**).

Finally, we asked if CD4+ and CCR4+ CD4+ T cells have increased proximity to phospho-STAT6+ tumour cells in STAT6^D419^-mutant samples. To do this, we measured the proportion of total tumour cells versus phospho-STAT6+ tumour cells that have direct cellular contact with CD4+/ CCR4+ CD4+ T cells (**Figure 6K**). When these measurements were performed, we found that the proportion of total tumour cells interacting with CD4+ T cells or CCR4+ CD4+ T cells was unchanged between STAT6^WT^ and STAT6^D419^-mutant tissues. However, we found that the proportion of CD4+ T cells interacting with phospho-STAT6+ tumour cells was significantly increased in STAT6^D419^-mutant biopsies, and the proportion of CCR4+ CD4+ T cells interacting with phospho-STAT6+ tumour cells trended towards increased in STAT6^D419^-mutant biopsies. Thus, these results strongly imply that the CCL17-CCR4 axis is over-activated in STAT6^D419^ rrDLCBL tissues, leading to alterations in TME organization, and favouring interactions between CD4+ T cells and phospho-STAT6+ tumour cells.

## Discussion

In the present study, we are the first to directly demonstrate that mutation of STAT6 at residue D419 is sufficient to drive microenvironmental remodelling, largely via the CCL17-CCR4 axis. Using an immunocompetent mouse model, we showed that STAT6^D419N^ mutant tumours have increased invasion of CCR4+ CD4+ T cells, coupled with resistance to doxorubicin treatment. Moreover, *ex vivo* functional assays demonstrated that STAT6^D419N^ tumour cells are directly chemoattractive to CCR4+ CD4+ T cells. When Eµ-Myc-mSTAT6^D419N^ tumour bearing mice were treated with the small molecule CCR4 inhibitor AZD2098, we observed that CD4+ T cell infiltration was reduced, and mice were re-sensitized to doxorubicin treatment. In human STAT^D419^ tumour samples, we confirmed that expression of phospho-STAT6 in tumour cells is increased relative to STAT6^WT^ rrDLBCL samples. As predicted by our mouse modelling, we observed increased tumour cell expression of CCL17 and increased cellular interactions of phospho-STAT6+ tumour cells with CD4+ T cells. Thus, our findings via mouse modelling are highly relevant to the human rrDLBCL population.

Given the increase in phospho-STAT6 expression observed in human and murine STAT6^D419^ DLBCL tumours (13), we initially hypothesized that STAT6^D419^-mutant tumour cells might be more chemoattractive to IL-4-producing CD4+ T cells, such as Th2 or Tfh cells, that may then function to maintain STAT6 phosphorylation. However, our data indicates that STAT6-mutant lymphoma tumour cells actually have an increased abundance of CCR4+ Th1-polarized CD4+ T cells, and not Th2 or Tfh cells. This is a surprising finding, especially given that CCR4 expression is typically associated with Th2 cells and Tregs; but we note that we are not the first to describe CCR4 expression on Th1 cells (22, 27, 28). Indeed, the m*STAT6*-mutant TME has increased Th1 associated cytokines, and m*STAT6* mutant tumour cells display evidence of an inflammatory gene signature. Concordant with previously published literature in DLBCL and other solid tumours, this inflammatory TME is associated with therapeutic resistance to doxorubicin (18–21, 29–31). Upon treatment of STAT6 mutant tumours with the CCR4 inhibitor AZD2098, CD4+ T cell invasion is reduced, and therapeutic sensitivity to doxorubicin is restored.

Our study utilized the power of multiplexed imaging to define tumor/ TME cellular heterogeneity within the context of spatial information in human rrDLBCL biopsies. We found that phospho-STAT6+ tumor cells were closely associated with CD4+ T cells, an interaction which when inhibited reversed doxorubicin-resistance. This emphasizes the importance of understanding not only what cell types are present in the TME, but also their association with each other. While flow cytometry allows screening of larger number of cells comprising the TME, PhenoCycler adds information regarding location. Of note, a current limitation of this study is the dependence upon region selection for the creation of a TMA and a single 2D section per tumor. However, a strength of this technology is the ability to detect cytokines/ chemokines and phospho-proteins in cells of the TME, which is notoriously difficult to do via flow cytometry. Indeed, in rrDLBCL patient samples, we found that phospho-STAT6 and CCL17 are more highly expressed in tumour cells of STAT6^D419^-mutant patients, and expression of phospho-STAT6 was positively correlated with tumour cell expression of CCL17 and CD4+ T cell expression of CCR4. These results expand our previous findings (14), to assert that phospho-STAT6 expression in rrDLBCL tumours contributes to TME remodelling via favouring interactions of tumour cells with the CD4 T cell compartment, largely due to the CCL17-CCR4 axis.

CCR4 has been established as a therapeutic target in Mycosis fungoides (MF) and Sezary syndrome (SS), due to tumour cell upregulation of CCR4. Indeed, the CCR4-targeted monoclonal antibody Mogamulizumab was FDA approved for the treatment of relapsed and refractory MF and SS in 2018 (32, 33). CCR4 has also been previously described as a potential therapeutic target in DLBCL. In a study using imaging mass cytometry (IMC), it was reported that CCR4 is expressed on PD1+ Tregs in the DLBCL and cHL TME. In DLBCL, CCR4+ Tregs are in close proximity to PD1+ CD4+ T cells and PD1+ CD8+ T cells, while these interactions are not present in cHL tumours (34). The authors suggest that these cellular interactions may be a mechanism of immune evasion in DLBCL relative to cHL and may be one factor that underlies the lack of therapeutic susceptibility of DLBCL tumours to immune checkpoint inhibitors. Accordingly, in a Phase I clinical trial (NCT03309878) Mogamulizumab was tested in combination with the anti-PD1 antibody Pembrolizumab for the treatment of rrDLBCL (35); however, due to the presence of serious adverse events, the trial was discontinued prior to Phase II. Our data support that *STAT6-* mutant rrDLBCL would benefit from Mogamulizumab, thus evaluating its efficacy as a monotherapy or finding a safe combination with other therapies may be warranted in future trials.

While our results indicate a critical role for the CCL17-CCR4 axis in STAT6^D419^-mediated microenvironmental remodelling, they do not preclude the potential for other mechanisms of action underlying STAT6^D419^-mediated doxorubicin resistance. For instance, STAT6 is a transcription factor that controls expression of genes such as CD23, CISH, IL2RA, BCL-6, BCL-XL, CDK2 (4, 6, 36) . These proteins play roles in B cell activation, resistance to apoptosis, cell cycling. Thus, future studies may investigate the cellular and molecular responses of STAT6^D419^ tumour cells to doxorubicin, with a specific focus on how altered regulation of these genes impacts tumour cell survival in an intact TME. As such, our study did not address specifically why CCR4+ CD4+ T cells engender therapeutic resistance in STAT6^D419N^ mutant tumours. In future studies, single cell transcriptomic approaches will be used to determine the specific phenotype of CCR4+ CD4+ T cells, and how they might drive tumour growth and chemoresistance.

In summary, our findings highlight the importance of investigations focused on the intricate interplay between DLBCL tumour cells and the TME. Indeed, the full functional impact of *STAT6^D419^* mutations could only be realized by utilizing an immunocompetent mouse model. Many gene mutations are predicted to modify the interaction of DLBCL with its TME, thus future studies should take a TME-centric approach to better understand how this relationship affects response to immune-based therapies, such as immunomodulatory agents, bi-specific T cell engagers and chimeric antigen receptor (CAR) T cells. Overall, a better understanding of the mechanisms of therapeutic resistance and immune escape in DLBCL could improve clinical outcomes, particularly for those with relapsed and refractory disease.

## Materials and Methods

All reagents and tools can be found in **Supplementary Table 3**.

### Cell Culture

MEF^ARF-/-^ and Phoenix-AMPHO cells were grown in DMEM media supplemented with 10% fetal bovine serum (FBS) and 0.5% penicillin-streptomycin (P/S). To generate mitotically inactive MEFs for co-culture, MEF^ARF-/-^ cells were given 40 Gy accumulated irradiation using a MultiRad225 (Faxitron) with a 0.5mm Cu filter and set to 225.0 kV and 17.8 mA. Following irradiation, cells were frozen in FBS supplemented with 10% DMSO. Eµ-Myc cells were co-cultured with irradiated MEF^ARF-/-^ (iMEF) cells in 50% DMEM/ 50% Iscove’s media, supplemented with FBS (10%), P/S (0.5%), and β-mercaptoethanol (50 nM).

To produce murine STAT6^WT^ and STAT6^D419N^ lymphoma cell lines, Phoenix-AMPHO packaging cells were transfected with pMIG plasmids containing WT or D419N mutant m*STAT6* and a GFP reporter. Retrovirus supernatant was used to transduce Eµ-Myc cells by spinoculation (centrifugation at 800 x g for 30 minutes at room temperature with 6 μg/mL polybrene, every 12 hours for 2 consecutive days). Successfully transduced GFP+ cells were sorted with a BD FACSAria Fusion Cell Sorter at the Lady Davis Institute Flow Cytometry Facility and were maintained in cell culture.

### Mouse Modelling

All mouse experiments were performed in accordance with an Animal Use Protocol approved by the McGill University Animal Care Committee. Mice were housed in the animal facility of the Lady Davis Institute for Medical Research.

1×10^6^ mSTAT6^WT^ or mSTAT6^D419N^ Eµ-Myc cells in 0.1mL of sterile PBS were injected into 8– 10-week-old female C57BL/6 mice via tail vein on Day 1. Tumour cells homed to lymph nodes, and disease burden was monitored using a VEVO-3100 Ultrasound machine, with Vevo LAB software used for tumour volume calculation. Mice were sacrificed if tumour volume in the cLN exceeded 100 mm^3^, or if they showed signs of weight loss and hind-leg paralysis.

Where indicated, mice were treated with doxorubicin (2 mg/mL; Jewish General Hospital Oncology Pharmacy) or AZD2098 (MedChemExpress). Doxorubicin was given as three doses of 3 mg/kg or one dose of 10 mg/kg, via intraperitoneal injection using a sterile 28-gauge needle. AZD2098 was purchased as powder, and was prepared at 2 mg/mL, in 10% DMSO and 90% corn oil. Mice were given twice weekly gavage, at a dose of 10 mg/kg At the indicated time points, mice were humanely sacrificed using isoflurane anesthesia, CO_2_, and cervical dislocation. Tumours were harvested and processed for flow cytometry, or were saved in FFPE blocks.

### Immunohistochemistry

For Ki67 staining, FFPE tumor and lymph node sections were deparaffinized using three changes of xylene and were hydrated through graded alcohols. Antigen-retrieval was performed for 20 minutes at 95 °C, using a PT Link Antigen Retrieval machine (Agilent) with Tris-EDTA pH 9.0 buffer. Endogenous peroxidases were quenched with 4.5% H_2_O_2_ for 15 minutes, and slides were blocked with 2% donkey serum for 30 minutes. They were then stained with Ki67 antibody (1:400; Cell Signaling Technology) for 30 minutes at 37°C and incubated in anti-rabbit secondary antibody for one hour at room temperature, followed by 1 minute DAB exposure. Slides were counterstained with hematoxylin, blued with blueing buffer, and were mounted with coverslips and Permount. For pSTAT6-CD4 co-staining, slides were prepared with the same deparaffinization, antigen retrieval, and quenching/blocking steps. Then slides were incubated with CD4 antibody (1:50; Invitrogen) for 30 minutes at 37°C and incubated in anti-rat secondary antibody for one hour at room temperature, followed by 2 minutes DAB exposure. Following DAB exposure, slides were again quenched and blocked. They were then incubated overnight at 4°C with pSTAT6 antibody (1:250; Cell Signaling Technology), and then were incubated in anti-rabbit secondary antibody for one hour, followed by a Magenta Red (Agilent) detection protocol. Slides were then counterstained with hematoxylin, blued with blueing buffer, and mounted. For all slides, images were acquired in brightfield with an AxioScan 7 (Zeiss) and were analyzed using QuPath 0.5.1-x64 software (37). Where applicable, staining was quantified based on percent positivity for a given marker, based on an intensity cut-off, or was quantified as H-Score. H-Score is calculated by indexing each cell within the tissue as having negative, weak, moderate, or strong staining for the given marker, based on three different intensity cut-offs. Then, the following formula was used to score the tissues, out of a maximum of 300: H-Score = (0* (% of negative cells)) + (1* (% of weak staining cells)) + (2* (% of moderate staining cells)) + (3* (% of strong staining cells))

### Flow Cytometry

cLN tumours were harvested at the indicated time points, and were gently crushed in a 1mL Eppendorf tube using a pestle. Following crushing, cells were passed through a 70 µm cell strainer. Single cells obtained from tumour dissociation were counted and 2×10^6^ cells per sample were plated in a 96-well V-bottom plate. Cells were incubated with fixable aqua live/dead stain (30 minutes; Invitrogen), Fc block (30 minutes), and then were incubated with fluorescently conjugated antibodies (30 minutes). For intracellular staining of transcription factors, cells were fixed and permeabilized (eBioscience), then were incubated in fluorescently conjugated intracellular staining antibodies diluted in permeabilization buffer. The flow antibodies used in this study can be found in **Supplementary Table 1**. Data was acquired on an LSRFortessa (BD; Lady Davis Institute Flow Cytometry Core), and FlowJo (BD) was used for all flow cytometry data analysis. Following analysis, the number of each cell type of interest was normalized to total tumour cell count, and data was presented as total number of each cell type found within each tumour.

### RNA Sequencing and qPCR

For RNA sequencing, CD19+ cells were purified from 3 biological replicates each of mSTAT6^WT^ and mSTAT6^D419N^ Eµ-Myc Day 11 cLN tumours, using CD19 microbeads (Miltenyi), as per manufacturer protocol. RNA was extracted from purified B cells using the Absolutely Total RNA Purification Kit (Agilent Technologies), as per manufacturer protocol. RNA sequencing was performed at Genome Quebec. Paired read 100 bp sequencing runs were performed on an Illumina NovaSeq 6000 S2 PE100.

For qPCR, RNA was extracted from tumour cells using the E.Z.N.A. total RNA isolation kit (OMEGA Bio-Tek). cDNA was synthesized from 1 mg of total RNA, using the iScript cDNA Synthesis Kit (Bio-Rad). mRNA expresson was quantified using the QuantStudio 7 Flex PCR System with SYBR Green.

### Ex Vivo Chemoattraction

For experiments with chemoattraction towards tumour cells *in vitro*, Eµ-Myc-mSTAT6^WT^ or Eµ-Myc-mSTAT6^D419N^ cells were plated at a confluence of 3×10^5^ cells/mL, overtop of iMEF cells. After 24 hours, tumour cells were stimulated with 5 ng/mL of murine (m) IL-4 (PeproTech) or vehicle control. After 1 hour, cells from a duplicate plate were collected for RNA isolation and qPCR analysis, to confirm CCL17 upregulation in mIL-4 stimulated Eµ-Myc-mSTAT6^D419N^ tumour cells. In tandem, spleens from female tumour naïve C57BL/6 mice were harvested and dissociated. Red blood cells (RBCs) were lysed for 10 minutes on ice (eBioscience). Splenocytes were treated with either 10 µM AZD2098 (MedChemExpress) or vehicle control for 1 hour at room temperature. Following tumour cell incubation in mIL-4 and splenocyte incubation in AZD2098, 1×10^6^ bulk splenocytes were plated overtop the stimulated or non-stimulated Eµ-Myc tumour cells, using a 5 µm pore transparent membrane. Splenocytes were allowed to migrate towards Eµ-Myc tumour cells for 16 hours. Migrated splenocytes were collected and stained with fixable aqua live/dead stain. They were then incubated with Fc block (30 minutes), then were stained with fluorescently conjugated antibodies to detect B cells, CD4+ T cells, CD8+ T cells, DCs, macrophages, NK cells, and eosinophils. Cells were not fixed, in order to preserve tumour cell endogenous GFP expression. The entire sample was acquired on an LSRFortessa, and FlowJo was used for analysis. The plated tumour cells were distinguished from migrated cells via expression of GFP and CD19, and tumour cells were excluded from the migrated cell counts.

For chemoattraction towards tumour supernatants, cLN tumours and spleens were harvested at day 11 post-Eµ-Myc-mSTAT6^WT^ and Eµ-Myc-mSTAT6^D419N^ tumour cell injection. cLNs were crushed in a 1mL Eppendorf tube with 500uL total volume of PBS and were incubated on ice for 20 minutes. Following incubation, tubes were spun down at 300 x g for 10 minutes, and tumour supernatants were collected (see below). Spleens were also dissociated, with 10-minute RBC lysis.

Following dissociation, CD4+ T cells were purified from splenocytes (Miltenyi), as per manufacturer protocol. Following purification, CD4+ T cells were treated for 1 hour at room temperature with either 10 µM AZD2098 or vehicle control. While CD4+ T cells were incubating in AZD2098, equal volumes of tumour supernatants or PBS control were plated in a 12-well plate. 5×10^5^ CD4+ T cells were plated overtop the Eµ-Myc-mSTAT6^WT^ and Eµ-Myc-mSTAT6^D419N^ tumour supernatants, using a 3 µm pore transparent membrane. CD4+ T cells were allowed to migrate towards tumour supernatants for 16 hours, then were collected for CD4+ T cell quantification and phenotyping by flow cytometry.

### Chemokine/Cytokine Profiling

At day 11 post-Eµ-Myc injection, Eµ-Myc-mSTAT6^WT^ and Eµ-Myc-mSTAT6^D419N^ tumours were dissociated, as described above. Tumour samples were allowed to incubate on ice on 20 minutes, then were spun down at 300 x g for 10 minutes. The supernatant was collected and was either used fresh for chemoattraction experiments or stored at -80 °C for cytokine profiling. The cell pellet was counted. To determine the cytokine/chemokine constitution of tumour supernatants, 44-Plex Mouse Cytokine/Chemokine profiling was performed with Eve Technologies (Calgary, AB, Canada), and cytokine/chemokine expression was normalized to total cell number in each tumour.

### PhenoCycler Staining

PhenoCycler staining was performed as described previously (24, 25, 38). FFPE lymphoma tissue sections were baked for 1 hour at 60 °C, deparaffinized using xylene, and were hydrated through graded alcohols. Antigen-retrieval was performed for 20 minutes at 95 °C, with Tris-EDTA pH 9.0 buffer. Tissue autofluorescence was quenched by bathing slides in a solution of 4.5% H_2_O_2_ and 20 mM NaOH prepared in PBS, sandwiched between LED lights at 25000 lux, for 45 minutes at room temperature. Slides were then washed with PBS (3 x 5 minutes) and hydration buffer (Akoya Biosciences; 2 x 5 minutes). Slides were equilibrated in staining buffer (Akoya Biosciences; 30 minutes) while the antibody staining cocktail was prepared in staining buffer supplemented with N, G, J, and S blockers (Akoya Biosciences) and oligonucleotide-conjugated antibodies diluted at the concentrations indicated in **Supplementary Table 2**. Antibody staining occurred in two steps. The first step used antibodies optimized for 30-minute incubation at 37 °C. After the first staining step, slides were washed in three changes of staining buffer before proceeding. The second step used antibodies optimized for overnight (ON) incubation at 4 °C. After overnight staining, slides were washed in three changes of staining buffer, and were fixed with 1.6% PFA for 10 minutes, ice-cold methanol for 5 minutes, and fixative solution (Akoya Biosciences) for 20 minutes. Slides were then stored until imaging with a PhenoCycler-Fusion instrument (up to 2 days).

The slides were mounted with a flow cell (Akoya Biosciences), to allow for automated fluidics that wash on and off fluorescently conjugated oligonucleotides that are complementary to the oligonucleotides that are conjugated to each antibody (aka “reporters”), while allowing the slide to stay mounted to a fluorescent microscope. To image slides, a reporter plate was prepared that included solutions containing PhenoCycler-Fusion buffer, DAPI, salmon sperm DNA, and up to three different reporters. The reporter plate solutions were washed onto the tissue cyclically, using the PhenoCycler-Fusion instrument, until all antibodies in the staining panel were visualized. The final outcome was qptiff files with highly multiplexed immunofluorescent images.

Following staining, the flow cell was removed using 24-hour incubation in Citrisolv. The Citrisolv was washed off, using grading alcohols, then H&E staining was performed. Slides were then mounted and imaged with an AxioScan 7 (Zeiss).

### PhenoCycler Data Analysis

Multiplexed qptiff images were uploaded to Enable Medicine, where cell segmentation was performed using DeepCell, with DRAQ5 as a nuclear marker and B-actin as a membrane marker. Quality control was used to exclude tumour cores of poor quality or individual cells that showed evidence of staining artifacts (ie. increased signal sum, decreased signal variation, and abnormalities in cell size or nuclear signal). Unsupervised clustering was then performed to identify major cell types, using Leiden clustering, set to detect 50 neighbours, with a UMAP minimum distance of 0.1 and a clustering resolution of 0.5. The accuracy of clustering was visually confirmed using Voronoi diagrams, and Leiden sub-clustering was used to further refine clusters to ensure accurate cell classification. Sub-clustering was also used to subclassify cells based on the expression of markers of interest. Spatial neighbour calculations were performed in Enable, by calculating the proportion of interacting cells for chosen cell types.

### Patient Data

The patient cohort used for PhenoCycler imaging consisted of FFPE preserved samples obtained from consented patients treated in Montreal QC, at the Jewish General Hospital. All biopsies were obtained in the context of routine clinical care. Biobanking of the patient plasma at the time of relapse and secondary use of plasma and FFPE samples for this project were approved by the research ethics board (REB 11-047, 15-036). Patients were all treated with curative intent R-CHOP-like regimens. Using banked specimens, one-millimeter cores of rrDLCBL FFPE tissue were arranged into TMA, at the Jewish General Hospital Research Pathology Core. 4uM TMA sections were mounted onto SuperFrost slides and were stained with a 52-plex PhenoCycler antibody cocktail, following the protocol described above. For STAT6^D419^ mutant samples, whole sections were mounted onto SuperFrost slides, with two sections from two different patients per slide.

Patient metadata included age, sex, cell-of-origin (COO), and STAT6 mutational status. Targeted sequencing and exome sequencing of the plasma and FFPE samples, to determine STAT6 mutational status, was previously reported by Rushton et al (39). Briefly, plasma samples were prepared into sequencing libraries using custom adaptors containing unique molecule identifiers. Libraries were enriched using a custom set of probes targeted to genes known to be involved in rrDLBCL pathology, including *STAT6,* and sequenced using 150-bp reads on Illumina MiSeq or HiSeq2500 platforms. FFPE samples were prepared and previously sequenced using whole exome capture. All sample somatic mutations were identified using Strelka2 and a custom bioinformatics pipeline previously described by Rushton et al (39).

### Statistical Analysis

A detailed list of statistics used, including test used, same size, and p-value, for each figure can be found in **Supplementary Table 4**.

### Data Availability Statement

All raw data will be made available upon reasonable request to the corresponding author.

