## Supplemental Material for "The CCL17-CCR4 axis is critical for mutant STAT6-mediated microenvironmental remodelling and therapeutic resistance in Relapsed/Refractory Diffuse Large B Cell Lymphoma"

**Proposed Manuscript Type:** Original Article

Madelyn J. Abraham[1,2], Cynthia Guilbert[1], Natascha Gagnon[1], Christophe Goncalves[1], Alexandre Benoit[1,2], Ryan Rys[1], Samuel E. J Preston[1,2], Ryan D. Morin[3], Wilson H. Miller Jr[1,2,4], Nathalie A. Johnson[1,2,4], Sonia V. del Rincon[1,2,4], and Koren K. Mann[1,2,4,5]

**Affiliations of Authors:** [1]Lady Davis Institute, Jewish General Hospital, Montreal, Quebec, Canada, [2]Division of Clinical and Translational Research, McGill University; Montreal, Quebec, Canada, [3]Department of Molecular Biology and Biochemistry, Simon Fraser University, Burnaby, British Columbia, Canada [4]Gerald Bronfman Department of Oncology, McGill University, Montreal, Quebec, Canada, [5]Department of Pharmacology and Therapeutics, McGill University, Montreal, Quebec, Canada.

**Running Title:** CCL17-CCR4 in STAT6-mutant rrDLBCL

### SUPPLEMENTARY FIGURES

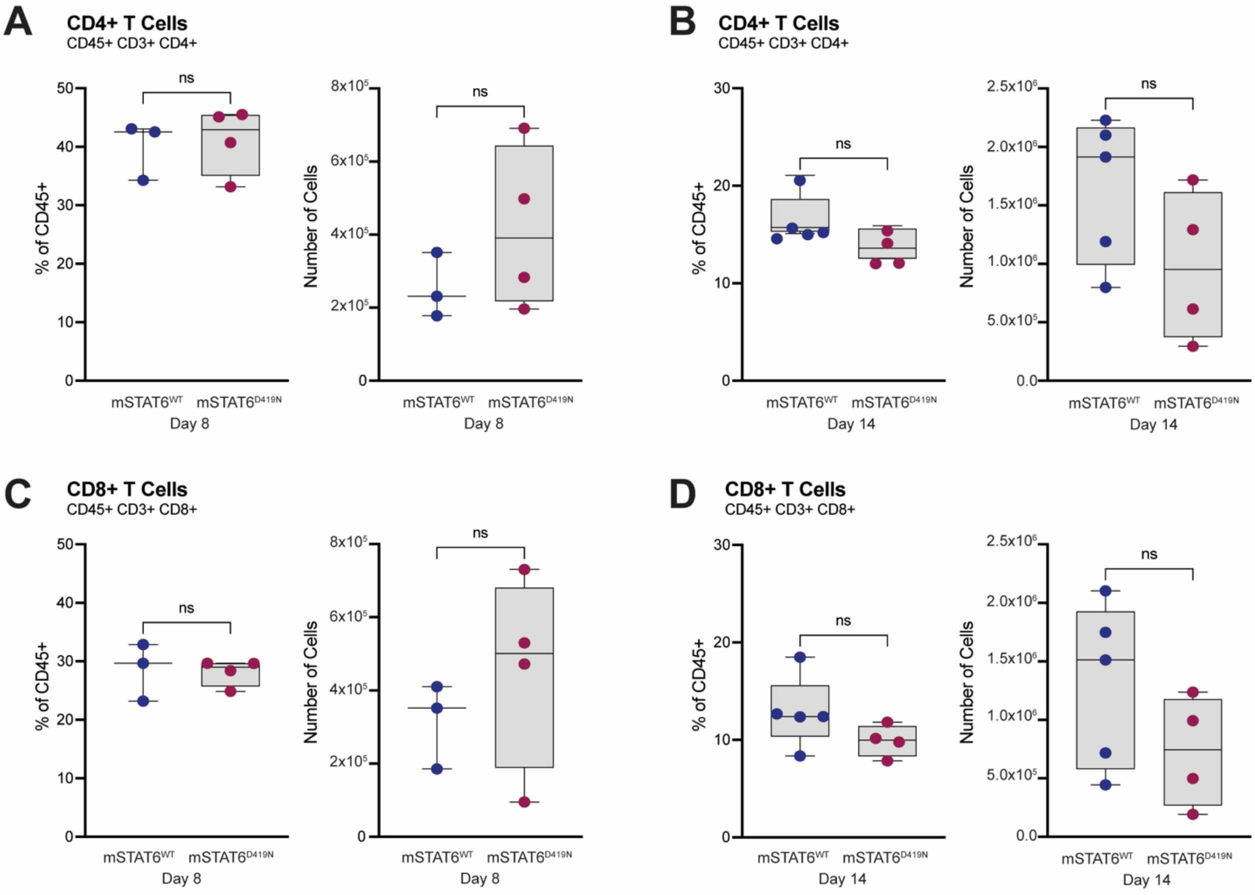


**Supplementary Figure 1: mSTAT6$^{D419N}$ Eμ-Myc tumours have no difference in number or proportion of CD4+ or CD8+ T cells at early and late disease as compared to mSTAT6$^{WT}$ Eμ-Myc tumours.**

**A.** Quantification of CD4+ T cells in day 8 mSTAT6$^{WT}$ and mSTAT6$^{D419N}$ Eμ-Myc tumours. CD4+ T cells are live, single cells, that are CD45+, CD3+, and CD4+. Data is expressed as a percentage of CD45+ cells in each tissue, and as the total number of CD4+ T cells in each tissue. **B.** Quantification of CD4+ T cells in day 14 mSTAT6$^{WT}$ and mSTAT6$^{D419N}$ Eμ-Myc tumours. **C.** Quantification of CD8+ T cells in day 8 mSTAT6$^{WT}$ and mSTAT6$^{D419N}$ Eμ-Myc tumours. CD8+ T cells are live, single cells, that are CD45+, CD3+, and CD8+. **D.** Quantification of CD8+ T cells in day 14 mSTAT6$^{WT}$ and mSTAT6$^{D419N}$ Eμ-Myc tumours.

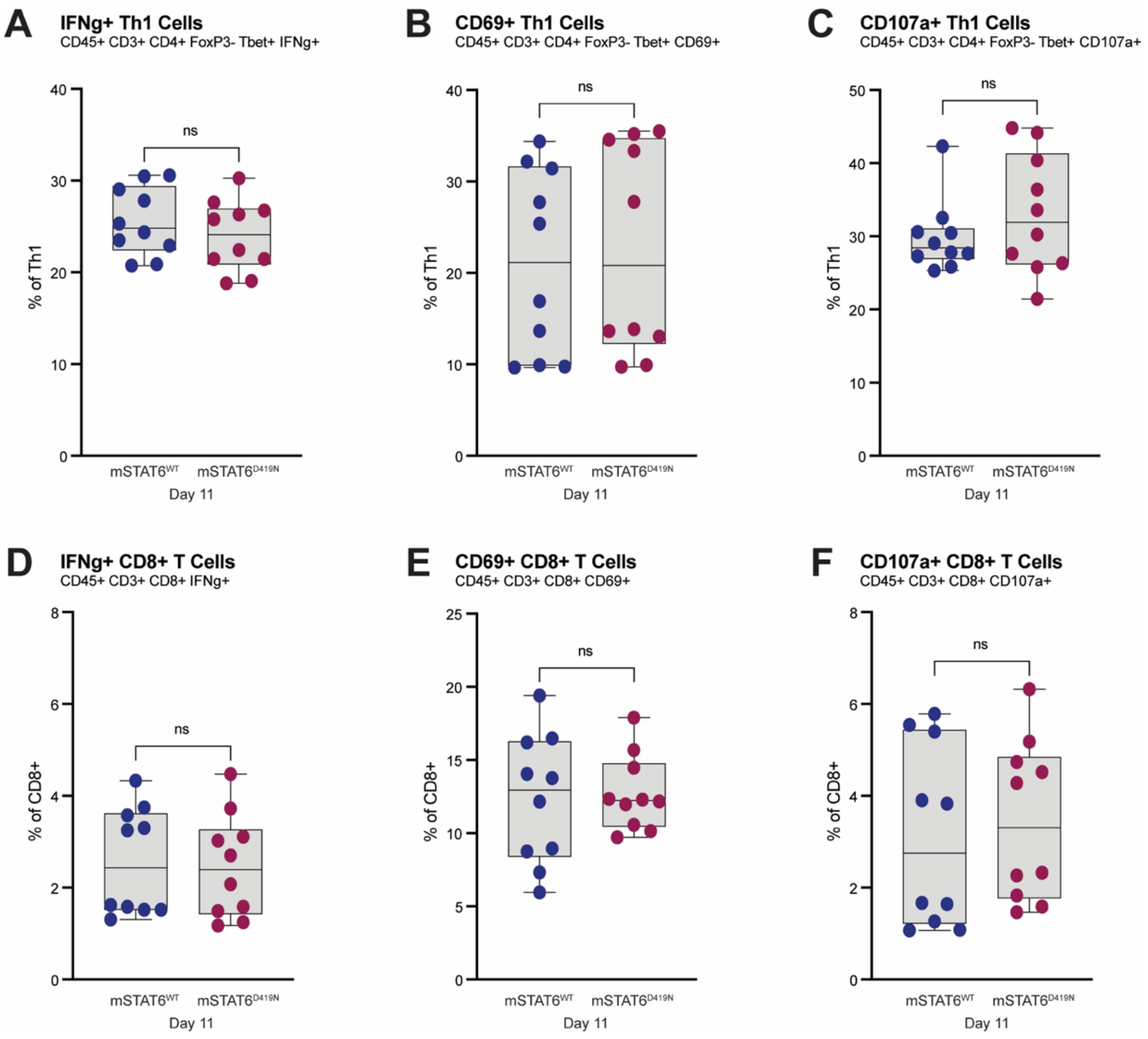


**Supplementary Figure 2: mSTAT6$^{D419N}$ Eμ-Myc tumours have no difference in expression of IFNγ, CD69, and CD107a in Th1 cells and CD8+ T cells as compared to mSTAT6$^{WT}$ Eμ-Myc tumours.**
**A.** Expression of IFNγ on Th1 cells in day 11 mSTAT6$^{WT}$ and mSTAT6$^{D419N}$ Eμ-Myc tumours, expressed as total percentage of Th1 cells that are IFNγ+ (CD45+ CD3+ CD4+ FoxP3- Tbet+ IFNγ+). **B.** Expression of CD69 on Th1 cells in day 11 mSTAT6$^{WT}$ and mSTAT6$^{D419N}$ Eμ-Myc tumours, expressed as total percentage of Th1 cells that are CD69+ (CD45+ CD3+ CD4+ FoxP3- Tbet+ CD69+). **C.** Expression of CD107a on Th1 cells in day 11 mSTAT6$^{WT}$ and mSTAT6$^{D419N}$ Eμ-Myc tumours, expressed as total percentage of Th1 cells that are CD107a+ (CD45+ CD3+ CD4+ FoxP3- Tbet+ CD107a+). **D.** Expression of IFNγ on CD8+ T cells in day 11 mSTAT6$^{WT}$ and mSTAT6$^{D419N}$ Eμ-Myc tumours, expressed as total percentage of CD8+ T cells that are IFNγ+ (CD45+ CD3+ CD8+ IFNγ+). **E.** Expression of CD69 on CD8+ T cells in day 11 mSTAT6$^{WT}$ and mSTAT6$^{D419N}$ Eμ-Myc tumours, expressed as total percentage of CD8+ T cells that are CD69+ (CD45+ CD3+ CD8+ CD69+). **F.** Expression of CD107a on CD8+ T cells in day 11 mSTAT6$^{WT}$ and mSTAT6$^{D419N}$ Eμ-Myc tumours, expressed as total percentage of CD8+ T cells that are CD107a+ (CD45+ CD3+ CD8+ CD107a+).

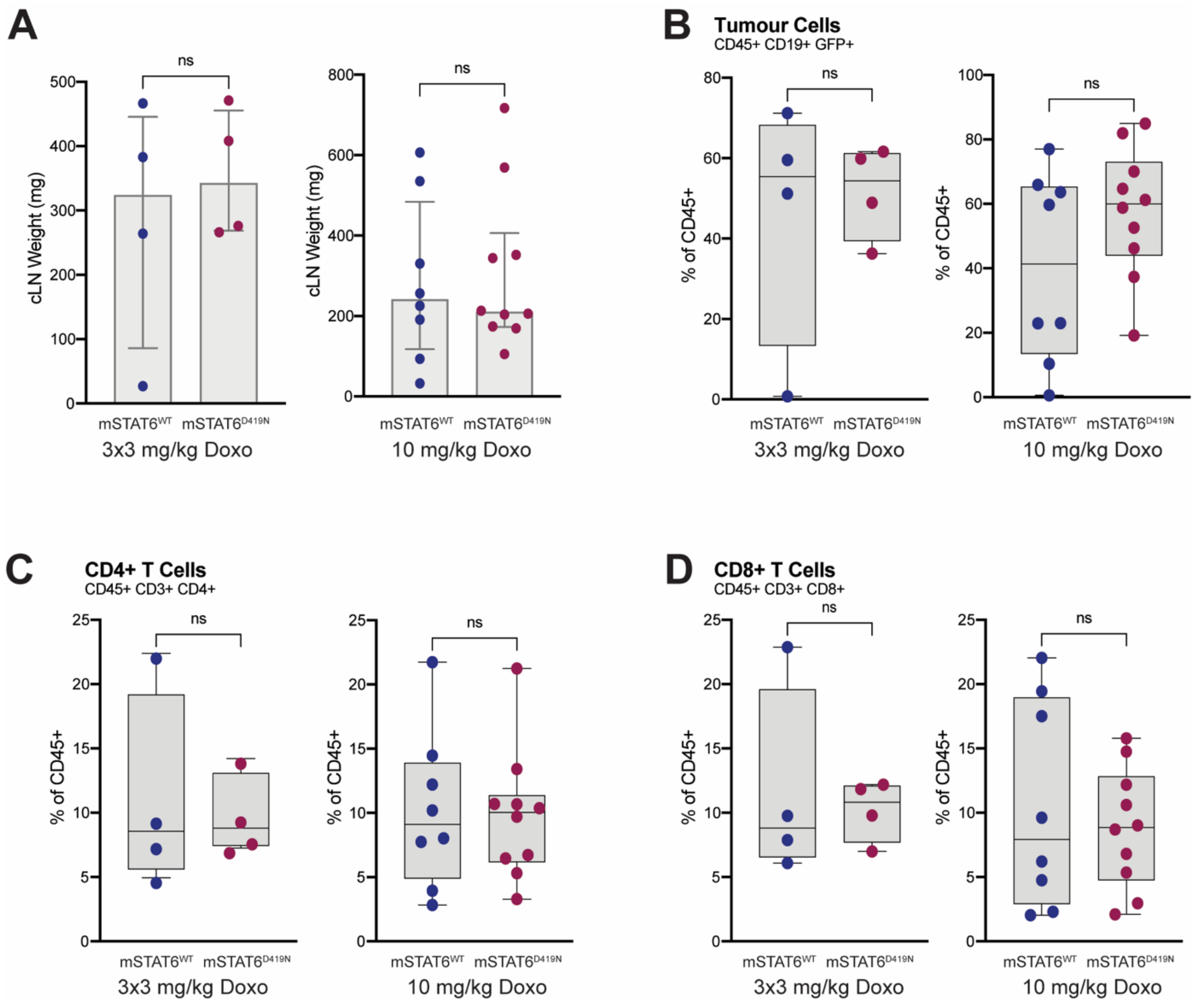


**Supplementary Figure 3: mSTAT6$^{WT}$ and mSTAT6$^{D419N}$ Eμ-Myc tumours have no difference in tumour burden or expression of CD4+ and CD8+ T cells, following relapse from doxorubicin treatment.**

**A.** Weights of mSTAT6$^{WT}$ and mSTAT6$^{D419N}$ relapsed Eμ-Myc tumours, following 3x3 mg/kg or 10mg/kg doxorubicin treatment. **B.** Quantification of tumour cells in relapsed mSTAT6$^{WT}$ and mSTAT6$^{D419N}$ Eμ-Myc tumours. Tumour cells are live, single cells, that are CD45+, CD19+, and GFP+. Data is expressed as a percentage of CD45+ cells in each tissue. **C.** Quantification of CD4+ T cells in relapsed mSTAT6$^{WT}$ and mSTAT6$^{D419N}$ Eμ-Myc tumours. CD4+ T cells are live, single cells, that are CD45+, CD3+, and CD4+. Data is expressed as a percentage of CD45+ cells in each tissue. **D.** Quantification of CD8+ T cells in relapsed mSTAT6$^{WT}$ and mSTAT6$^{D419N}$ Eμ-Myc tumours. CD8+ T cells are live, single cells, that are CD45+, CD3+, and CD8+. Data is expressed as a percentage of CD45+ cells in each tissue.

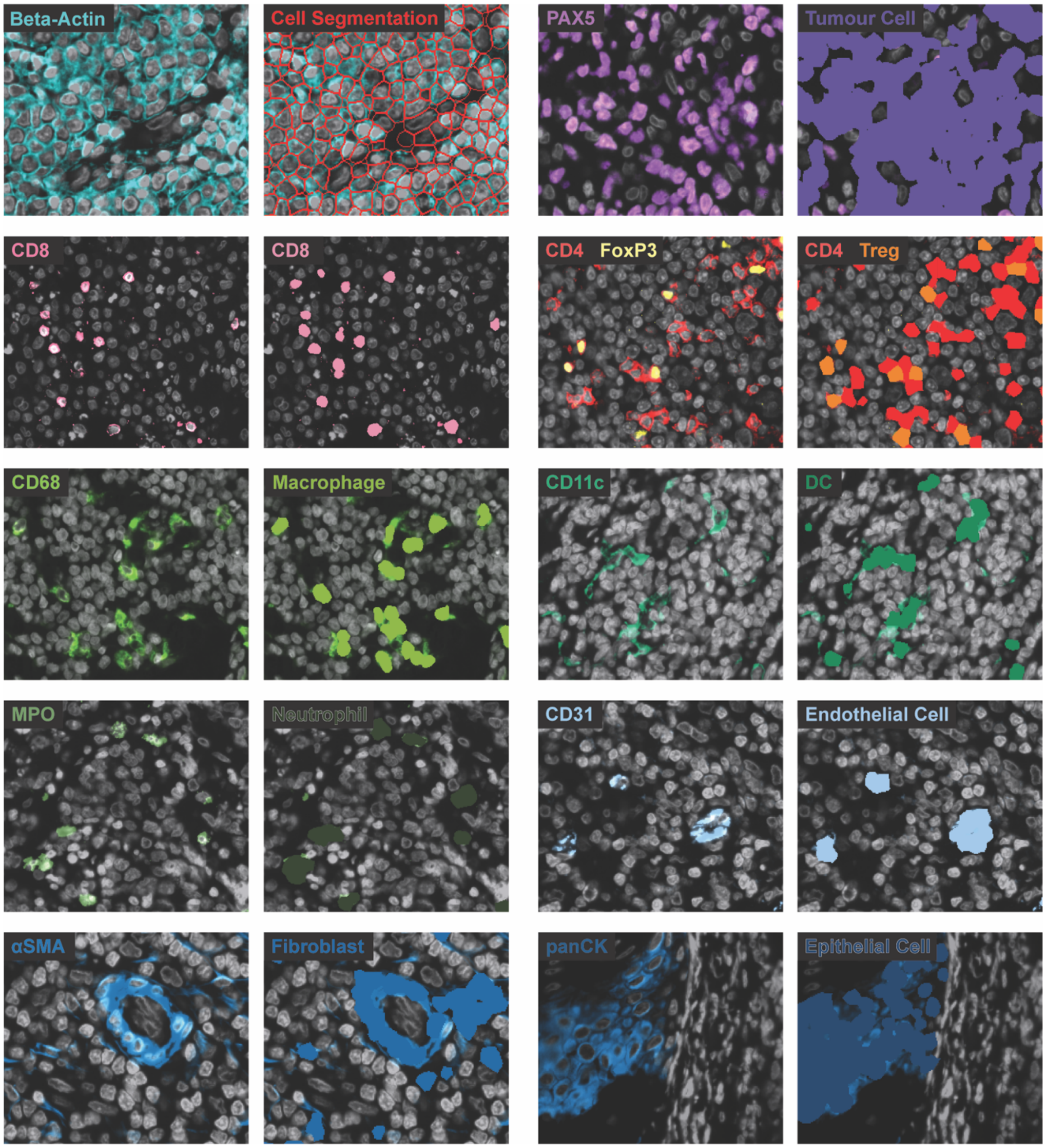


**Supplementary Figure 4: Cell segmentation and classification of human DLBCL PhenoCycler images.**

Cell segmentation overlay and Voronoi overlays showing accurate classification of different cell types within human lymphoma tissues.

#### SUPPLEMENTARY TABLES

**Supplementary Table 1. Antibodies for Flow Cytometry**

| Antibody | Clone | Fluorophore | Supplier | Catalog No. |
|---|---|---|---|---|
| CD45 | 30-F11 | BUV395 | eBioscience | 363-0451-82 |
| CD19 | 1D3 | AF700 | BD Pharmingen | 557958 |
| CD19 | 1D3 | PE-Cy7 | BD Pharmingen | 552854 |
| CD3 | 145-2C11 | BV650 | BD Horizon | 564378 |
| CD4 | RM4-5 | APC-Cy7 | BD Pharmingen | 565650 |
| CD8 | 53-6.7 | PerCP-Cy5.5 | BD Pharmingen | 551162 |
| PD-L1 | MIH5 | BUV737 | BD OptiBuild | 741877 |
| CXCR5 | L138D7 | PE/Dazzle594 | Biolegend | 145522 |
| PD-1 | J43 | BV421 | BD Horizon | 562584 |
| GATA3 | L50-823 | BV711 | BD Horizon | 565449 |
| FoxP3 | FJK-16s | FITC | Invitrogen | 11-5773-82 |
| Tbet | O4-46 | AF647 | BD Pharmingen | 561267 |
| CCR4 | 2G12 | PE-Cy7 | Biolegend | 131214 |
| IFNγ | XMG1.2 | PE | BD Pharmingen | 562020 |
| CD69 | H1.2F3 | PE-Cy5 | Biolegend | 104510 |
| CD107a | 1D4B | BV786 | BD Horizon | 564349 |
| CD11b | M1/70 | e450 | Invitrogen | 48-0112-82 |
| CD11c | HL3 | BV786 | BD Horizon | 563735 |
| F4/80 | BM8 | PE | Invitrogen | 12-4801-82 |
| NKp46 | 29A1.4 | APC | eBioscience | 11-3351-80 |
| SiglecF | E50-2440 | PE-CF594 | BD Horizon | 562757 |

**Supplementary Table 2: Antibodies for PhenoCycler Staining (52-plex)**

| Antibody | Barcode | Fluorophore | Staining Concentration |
|---|---|---|---|
| CD31 | BX001 | AF750 | 1 in 100, ON |
| TIGIT | BX002 | ATTO550 | 1 in 100, ON |
| CD4 | BX003 | AF647 | 1 in 200, ON |
| pSTAT6 | BX004 | AF750 | 1 in 25, ON |
| TIM3 | BX005 | ATTO550 | 1 in 200, ON |
| TCF1/7 | BX006 | AF647 | 1 in 100, ON |
| CD20 | BX007 | AF750 | 1 in 400, ON |
| CD15 | BX010 | AF647 | 1 in 100, ON |
| αSMA | BX013 | AF750 | 1 in 200, ON |
| PAX5 | BX014 | ATTO550 | 1 in 200, ON |
| CD68 | BX015 | AF647 | 1 in 200, ON |
| CCL17 | BX016 | AF647 | 1 in 100, ON |
| CD45RO | BX017 | ATTO550 | 1 in 200, ON |
| panCK | BX019 | AF750 | 1 in 200, ON |
| IFNγ | BX020 | AF647 | 1 in 200, ON |
| CD45 | BX021 | AF647 | 1 in 200, ON |
| CCL22 | BX022 | AF750 | 1 in 100, ON |
| NaK-ATPase | BX023 | ATTO550 | 1 in 100, ON |
| CD11c | BX024 | AF647 | 1 in 100, 30 min 37 °C |
| CD8 | BX026 | ATTO550 | 1 in 200, ON |
| CX3CR1 | BX027 | AF647 | 1 in 100, ON |
| CD56 | BX028 | ATTO550 | 1 in 200, ON |
| HLA-A | BX029 | ATTO550 | 1 in 200, ON |
| FoxP3 | BX031 | AF647 | 1 in 200, ON |
| CD21 | BX032 | ATTO550 | 1 in 200, ON |
| HLA-DR | BX033 | AF647 | 1 in 400, ON |
| CD11b | BX034 | ATTO550 | 1 in 200, ON |
| CD163 | BX035 | ATTO550 | 1 in 200, ON |
| CTLA4 | BX036 | AF647 | 1 in 100, ON |
| CD14 | BX037 | ATTO550 | 1 in 200, ON |
| VISTA | BX040 | ATTO550 | 1 in 100, ON |
| GZMB | BX041 | ATTO550 | 1 in 200, ON |
| CX3CL1 | BX042 | AF647 | 1 in 100, 30 min 37 °C |
| CD19 | BX043 | ATTO550 | 1 in 200, ON |
| MNK1 | BX045 | AF647 | 1 in 50, ON |
| PD-1 | BX046 | AF647 | 1 in 100, ON |
| Ki67 | BX047 | ATTO550 | 1 in 200, ON |
| GATA3 | BX049 | ATTO550 | 1 in 100, ON |
| CD86 | BX050 | AF647 | 1 in 100, ON |
| Tbet | BX052 | ATTO550 | 1 in 100, ON |
| p-eIF4E | BX054 | AF647 | 1 in 50, 30 min 37 °C |
| LAG3 | BX055 | AF647 | 1 in 100, ON |

| | | | |
|---|---|---|---|
| TOX | BX060 | ATTO550 | 1 in 200, ON |
| PD-L1 | BX067 | AF647 | 1 in 200, ON |
| CitH3 | BX078 | ATTO550 | 1 in 100, ON |
| CD3 | BX080 | AF647 | 1 in 200, ON |
| CD38 | BX089 | ATTO550 | 1 in 100, ON |
| MPO | BX098 | ATTO550 | 1 in 200, ON |
| CCR4 | BX106 | AF647 | 1 in 100, ON |
| CD45RA | BX113 | ATTO550 | 1 in 100, ON |
| B-Actin | BX117 | AF750 | 1 in 200, ON |
| eIF4E | BX518 | AF647 | 1 in 100, ON |

**Supplementary Table 3: Reagents and Tools**

| Reagent or resource | Reference or source | Identifier |
|---|---|---|
| **Cell lines** | | |
| Eμ-Myc Parental | Dr. Jerry Pelletier | NA |
| $MEF^{ARF-/-}$ | Dr. Jerry Pelletier | NA |
| Phoenix-AMPHO | ATCC | CRL-3212 |
| Eμ-Myc $mSTAT6^{WT}$ | This study | NA |
| Eμ-Myc $mSTAT6^{D419N}$ | This study | NA |
| **Antibodies for IHC** | | |
| Ki67 Rabbit mAb | Cell Signaling Technology | 12202S |
| CD4 Monoclonal Antibody | Invitrogen | 14-0042-82 |
| Phospho-STAT6 (Tyr641) Rabbit mAb | Cell Signaling Technology | 56554S |
| EnVision+ System- HRP Labellled Polymer Anti-Rabbit | Dako | K4003 |
| Goat Anti-Rat IgG H+L (HRP Polymer) | Abcam | ab214882 |
| **Dyes and Stains** | | |
| LIVE/DEAD Fixable Aqua Dead Cell Stain Kit | ThermoFisher | L34966 |
| DAPI | ThermoFisher | D1306 |
| DRAQ5 | Biolegend | 424101 |
| Harris' Alum Hematoxylin Mercury Free | Sigma-Aldrich | 638A-85 |
| Eosin Y Solution | Sigma-Aldrich | HT110116-500ML |
| ImmPACT DAB Substrate Kit, Peroxidase | Vector Laboratories | SK-4105 |
| EnVision FLEX HRP Magenta Substrate Chromogen System | Agilient Technologies | GV92511-2 |
| **Drugs, reagents, and other consumables** | | |
| Doxorubicin | Jewish General Hospital Oncology Pharmacy | 1201 |
| AZD2098 | MedChemExpress | HY-U00064 |
| Recombinant Murine IL-4 | PeproTech | 214-14 |
| FoxP3/ Transcription Factor Fixation/ Permeabilization Concentrate and Diluent | eBioscience | 00-5521-00 |
| CD19 MicroBeads, mouse | Miltenyi Biotec | 130-121-301 |
| CD4+ T Cell Isolation Kit, mouse | Miltenyi Biotec | 130-104-454 |
| LS Columns | Miltenyi Biotec | 130-042-401 |
| 5.0 um 6-well PET insert | Sarstedt | 83.3930.500 |
| 3.0 um 12-well PET insert | Sarstedt | 83.3931.300 |
| Sample Kit for PhenoCycler-Fusion | Akoya Biosciences | 7000017 |
| 10X Buffer Kit for PhenoCycler-Fusion | Akoya Biosciences | 7000019 |
| Antibody Conjugation Kit | Akoya Biosciences | 7000009 |

| | | |
|---|---|---|
| Salmon sperm DNA, sheared (10mg/mL) | Invitrogen | AM9680 |
| Permount Mounting Medium | Fisher Scientific | SP15-500 |
| Trypan Blue, 0.4% Solution | Wisent | 609-130-EL |
| RPMI | Wisent | 350-000-CL |
| ISCOVE | Wisent | 319-105-CL |
| DMEM | Wisent | 319-005-CL |
| Fetal bovine serum | Wisent | 080-450 |
| Penicillin-streptomycin | Wisent | 450-201-EL |
| β-mercaptoethanol | Sigma | M3148-100mL |
| **Software** | | |
| FlowJo v10.10 | BD Biosciences | |
| Prism (version 6.0) | GraphPad | |
| Illustrator 2024 | Adobe | |
| QuPath v0.5.1 | | |
| Enable Medicine Platform | Enable Medicine | |
| **Instruments** | | |
| BD LSRFortessa Flow Cytometer | BD Bioscience | |
| FACSAria Fusion Cell Sorter | BD Bioscience | |
| VEVO-3100 | FUJIFILM VisualSonics | |
| Mastercycler X50a | Eppendorf | |
| QuantStudio 7 Flex | Applied Biosystems by Life Technologies | |
| PT Link, Pre-Treatment Module for Tissue Specimens | Agilent Dako | |
| AxioScan 7 Microscope Slide Scanner | Zeiss | |
| Countess 3 Automated Cell Counter | ThermoFisher | AMQAX2000 |
| PhenoCycler-Fusion | Akoya Bioscience | |

**Supplementary Table 4: Statistics**

| Figure | Test Used | n | Comparisons and p-values |
|---|---|---|---|
| **Figure 1** | | | |
| 1C | 2way ANOVA, with Sidak's multiple comparison test | mSTAT6$^{WT}$ = 5<br>mSTAT6$^{D419N}$ = 5 | Day 4 mSTAT6$^{WT}$ vs Day 4 mSTAT6$^{D419N}$, p = 0.5284<br>Day 8 mSTAT6$^{WT}$ vs Day 8 mSTAT6$^{D419N}$, p = 2372<br>Day 11 mSTAT6$^{WT}$ vs Day 11 mSTAT6$^{D419N}$, p = 0.0217<br>Day 13 mSTAT6$^{WT}$ vs Day 13 mSTAT6$^{D419N}$, p = 0.5142 |
| 1E | 2way ANOVA, with uncorrected Fisher's LSD | D8 mSTAT6$^{WT}$ = 5<br>D8 mSTAT6$^{D419N}$ = 5<br>D11 mSTAT6$^{WT}$ = 8<br>D11 mSTAT6$^{D419N}$ = 8<br>D14 mSTAT6$^{WT}$ = 5<br>D14 mSTAT6$^{D419N}$ = 5 | Day 8 mSTAT6$^{WT}$ vs Day 8 mSTAT6$^{D419N}$, p = 0.7096<br>Day 11 mSTAT6$^{WT}$ vs Day 11 mSTAT6$^{D419N}$, p = 0.0001<br>Day 14 mSTAT6$^{WT}$ vs Day 14 mSTAT6$^{D419N}$, p = 0.8429 |
| 1F | Unpaired t test, two-tailed | D11 mSTAT6$^{WT}$ = 8<br>D11 mSTAT6$^{D419N}$ = 8 | Day 11 mSTAT6$^{WT}$ vs Day 11 mSTAT6$^{D419N}$, p = 0.8012 |
| 1H | Unpaired t test, two-tailed | D11 mSTAT6$^{WT}$ = 8<br>D11 mSTAT6$^{D419N}$ = 8 | CD4 H-score: Day 11 mSTAT6$^{WT}$ vs Day 11 mSTAT6$^{D419N}$, p = 0.0047<br>Phospho-STAT6 H-score: Day 11 mSTAT6$^{WT}$ vs Day 11 mSTAT6$^{D419N}$, p = 0.0138 |
| 1I | Pearson correlation | Number of pairs = 16<br>D11 mSTAT6$^{WT}$ = 8<br>D11 mSTAT6$^{D419N}$ = 8 | r = 0.7610<br>p = 0.0006 |
| 1J | Unpaired t test, two-tailed | D11 mSTAT6$^{WT}$ = 8<br>D11 mSTAT6$^{D419N}$ = 8 | % of CD45: Day 11 mSTAT6$^{WT}$ vs Day 11 mSTAT6$^{D419N}$, p = 0.0047<br>Number of cells: Day 11 mSTAT6$^{WT}$ vs Day 11 mSTAT6$^{D419N}$, p = 0.0267 |
| 1K | Unpaired t test, two-tailed | D11 mSTAT6$^{WT}$ = 8<br>D11 mSTAT6$^{D419N}$ = 8 | % of CD45: Day 11 mSTAT6$^{WT}$ vs Day 11 mSTAT6$^{D419N}$, p = 0.3459<br>Number of cells: Day 11 mSTAT6$^{WT}$ vs Day 11 mSTAT6$^{D419N}$, p = 0.4188 |
| **Figure 2** | | | |
| 2A | Unpaired t test, two-tailed | D11 mSTAT6$^{WT}$ = 10<br>D11 mSTAT6$^{D419N}$ = 10 | Day 11 mSTAT6$^{WT}$ vs Day 11 mSTAT6$^{D419N}$, p = 0.0122 |
| 2B | Unpaired t test, two-tailed | Tregs/ Tfh D11 mSTAT6$^{WT}$ = 7<br>Tregs/ Tfh D11 mSTAT6$^{D419N}$ = 7<br>Th1/ Th2 D11 mSTAT6$^{WT}$ = 10<br>Th1/ Th2 D11 mSTAT6$^{D419N}$ = 10 | Tregs: mSTAT6$^{WT}$ vs mSTAT6$^{D419N}$, p = 0.0793<br>Tfh: mSTAT6$^{WT}$ vs mSTAT6$^{D419N}$, p = 0.9008<br>Th1: mSTAT6$^{WT}$ vs mSTAT6$^{D419N}$, p = 0.0153<br>Th2: mSTAT6$^{WT}$ vs mSTAT6$^{D419N}$, p = 0.0943 |
| 2C | Unpaired t test, two-tailed | D11 mSTAT6$^{WT}$ = 10<br>D11 mSTAT6$^{D419N}$ = 10 | CCR4+ Th1: mSTAT6$^{WT}$ vs mSTAT6$^{D419N}$, p = 0.0317 |

| | | | |
|---|---|---|---|
| | | | CCR4+ MFI Th1: mSTAT6$^{WT}$ vs mSTAT6$^{D419N}$, p = 0.0017 |
| **Figure 3** | | | |
| 3B | 2way ANOVA, with Tukey's multiple comparison test | mSTAT6$^{WT}$ -IL-4 = 3<br>mSTAT6$^{WT}$ +IL-4 = 5<br>mSTAT6$^{D419N}$ -IL-4 = 3<br>mSTAT6$^{D419N}$ +IL-4 = 5 | mSTAT6$^{WT}$ -IL-4 vs mSTAT6$^{WT}$ +IL-4, p = 0.7936<br>mSTAT6$^{D419N}$ -IL-4 vs mSTAT6$^{D419N}$ +IL-4, p = 0.0053 |
| 3C | 2way ANOVA, with Tukey's multiple comparison test | mSTAT6$^{WT}$ -IL-4 = 5<br>mSTAT6$^{WT}$ +IL-4 = 5<br>mSTAT6$^{WT}$ +IL-4 +AZD2098 = 5<br>mSTAT6$^{D419N}$ -IL-4 = 5<br>mSTAT6$^{D419N}$ +IL-4 = 5<br>mSTAT6$^{D419N}$ +IL-4 +AZD2098 = 5 | CD4: mSTAT6$^{WT}$ -IL-4 vs mSTAT6$^{WT}$ +IL-4, p = 0.0008<br>CD4: mSTAT6$^{WT}$ +IL-4 vs mSTAT6$^{WT}$ +IL-4 +AZD2098, p < 0.0001<br>CD4: mSTAT6$^{WT}$ +IL-4 vs mSTAT6$^{D419N}$ +IL-4, p < 0.0001<br>CD4: mSTAT6$^{D419N}$ -IL-4 vs mSTAT6$^{D419N}$ +IL-4, p < 0.0001<br>CD4: mSTAT6$^{D419N}$ +IL-4 vs mSTAT6$^{D419N}$ +IL-4 +AZD2098, p < 0.0001<br>CD8, NK cell, macrophage, DC, eosinophil: mSTAT6$^{WT}$ -IL-4 vs mSTAT6$^{WT}$ +IL-4, p > 0.9999<br>CD8, NK cell, macrophage, DC, eosinophil: mSTAT6$^{WT}$ +IL-4 vs mSTAT6$^{WT}$ +IL-4 +AZD2098, p > 0.9999<br>CD8, NK cell, macrophage, DC, eosinophil: mSTAT6$^{WT}$ +IL-4 vs mSTAT6$^{D419N}$ +IL-4, p > 0.9999<br>CD8, NK cell, macrophage, DC, eosinophil: mSTAT6$^{D419N}$ -IL-4 vs mSTAT6$^{D419N}$ +IL-4, p > 0.9999<br>CD8, NK cell, macrophage, DC, eosinophil: mSTAT6$^{D419N}$ +IL-4 vs mSTAT6$^{D419N}$ +IL-4 +AZD2098, p > 0.9999 |
| **Figure 4** | | | |
| 4B | 2way ANOVA, with Sidak's multiple comparison test | mSTAT6$^{WT}$ +Veh = 5<br>mSTAT6$^{D419N}$+Veh = 5<br>mSTAT6$^{WT}$ + 3x3 mg/kg Doxo = 4<br>mSTAT6$^{D419N}$ + 3x3 mg/kg Doxo = 5 | mSTAT6$^{WT}$ +Veh vs mSTAT6$^{D419N}$+Veh, p = 0.8641<br>mSTAT6$^{WT}$ +Veh vs mSTAT6$^{WT}$ + 3x3 mg/kg Doxo, p = 0.0009<br>mSTAT6$^{D419N}$+Veh vs mSTAT6$^{D419N}$ + 3x3 mg/kg Doxo, p = 0.8151<br>mSTAT6$^{WT}$ + 3x3 mg/kg Doxo vs mSTAT6$^{D419N}$ + 3x3 mg/kg Doxo, p = 0.0511 |
| 4D | Gehan-Breslow-Wilcoxon test | mSTAT6$^{WT}$ + 3x3 mg/kg Doxo = 10<br>mSTAT6$^{D419N}$ + 3x3 mg/kg Doxo = 10 | mSTAT6$^{WT}$ + 3x3 mg/kg Doxo vs mSTAT6$^{D419N}$ + 3x3 mg/kg Doxo, p = 0.0764 |

| | | | |
|---|---|---|---|
| 4F | 2way ANOVA, with uncorrected Fisher's LSD | $mSTAT6^{WT}$ + 10 mg/kg Doxo = 10<br>$mSTAT6^{D419N}$ + 10 mg/kg Doxo = 10 | Pre: $mSTAT6^{WT}$ + 10 mg/kg Doxo vs $mSTAT6^{D419N}$ + 10 mg/kg Doxo, p = 0.2061<br>Post: $mSTAT6^{WT}$ + 10 mg/kg Doxo vs $mSTAT6^{D419N}$ + 10 mg/kg Doxo, p = 0.0023<br>$mSTAT6^{WT}$: Pre vs Post, p = 0.0615<br>$mSTAT6^{D419N}$: Pre vs Post, p = 0.0164 |
| 4H | Gehan-Breslow-Wilcoxon test | $mSTAT6^{WT}$ + 10 mg/kg Doxo = 10<br>$mSTAT6^{D419N}$ + 10 mg/kg Doxo = 10 | $mSTAT6^{WT}$ + 10 mg/kg Doxo vs $mSTAT6^{D419N}$ + 10 mg/kg Doxo, p = 0.1211 |
| **Figure 5** | | | |
| 5B | 2way ANOVA, with Tukey's multiple comparison test | $mSTAT6^{WT}$ + Veh= 9<br>$mSTAT6^{D419N}$ + Veh = 9<br>$mSTAT6^{WT}$ + 10 mg/kg AZD2098 = 9<br>$mSTAT6^{D419N}$ + 10 mg/kg AZD2098 = 8 | $mSTAT6^{WT}$ + Veh vs $mSTAT6^{D419N}$ + Veh, p = 0.0774<br>$mSTAT6^{WT}$ + Veh vs $mSTAT6^{WT}$ + 10 mg/kg AZD2098, p = 0.9613<br>$mSTAT6^{D419N}$ + Veh vs $mSTAT6^{D419N}$ + 10 mg/kg AZD2098, p = 0.0445 |
| 5C | 2way ANOVA, with Tukey's multiple comparison test | $mSTAT6^{WT}$ + Veh= 9<br>$mSTAT6^{D419N}$ + Veh = 9<br>$mSTAT6^{WT}$ + 10 mg/kg AZD2098 = 9<br>$mSTAT6^{D419N}$ + 10 mg/kg AZD2098 = 8 | Treg: $mSTAT6^{WT}$ + Veh vs $mSTAT6^{D419N}$ + Veh, p = 0.9940<br>Treg: $mSTAT6^{WT}$ + Veh vs $mSTAT6^{WT}$ + 10 mg/kg AZD2098, p = 0.6536<br>Treg: $mSTAT6^{D419N}$ + Veh vs $mSTAT6^{D419N}$ + 10 mg/kg AZD2098, p = 0.3958<br>Tfh: $mSTAT6^{WT}$ + Veh vs $mSTAT6^{D419N}$ + Veh, p = 0.7400<br>Tfh: $mSTAT6^{WT}$ + Veh vs $mSTAT6^{WT}$ + 10 mg/kg AZD2098, p = 0.9737<br>Tfh: $mSTAT6^{D419N}$ + Veh vs $mSTAT6^{D419N}$ + 10 mg/kg AZD2098, p = 0.9185<br>Th1: $mSTAT6^{WT}$ + Veh vs $mSTAT6^{D419N}$ + Veh, p = 0.0551<br>Th1: $mSTAT6^{WT}$ + Veh vs $mSTAT6^{WT}$ + 10 mg/kg AZD2098, p = 0.9970<br>Th1: $mSTAT6^{D419N}$ + Veh vs $mSTAT6^{D419N}$ + 10 mg/kg AZD2098, p = 0.1324<br>Th2: $mSTAT6^{WT}$ + Veh vs $mSTAT6^{D419N}$ + Veh, p = 0.7550<br>Th2: $mSTAT6^{WT}$ + Veh vs $mSTAT6^{WT}$ + 10 mg/kg AZD2098, p = 0.1055<br>Th2: $mSTAT6^{D419N}$ + Veh vs $mSTAT6^{D419N}$ + 10 mg/kg AZD2098, p = 0.0170 |
| 5D | 3way ANOVA | $mSTAT6^{WT}$ + Veh= 3<br>$mSTAT6^{D419N}$ + Veh =4 | Time, p < 0.0001<br>$mSTAT6^{WT}$ vs $mSTAT6^{D419N}$, p = 0.8843 |

| | | | |
|---|---|---|---|
| | | $mSTAT6^{WT}$ + 10 mg/kg AZD2098 = 3<br>$mSTAT6^{D419N}$ + 10 mg/kg AZD2098 = 5 | Veh vs AZD2098, p = 0.3360 |
| 5F | 2way ANOVA, with uncorrected Fisher's LSD | $mSTAT6^{WT}$ + Doxo = 5<br>$mSTAT6^{D419N}$ + Doxo = 5<br>$mSTAT6^{WT}$ + Doxo + AZD2098 = 5<br>$mSTAT6^{D419N}$ + Doxo + AZD2098 = 5 | Pre: $mSTAT6^{WT}$ + Doxo vs $mSTAT6^{D419N}$ + Doxo, p = 0.6611<br>Pre: $mSTAT6^{WT}$ + Doxo vs $mSTAT6^{WT}$ + Doxo + AZD2098, p = 0.7703<br>Pre: $mSTAT6^{D419N}$ + Doxo vs $mSTAT6^{D419N}$ + Doxo + AZD2098, p = 0.4867<br>Post: $mSTAT6^{WT}$ + Doxo vs $mSTAT6^{D419N}$ + Doxo, p = 0.0003<br>Post: $mSTAT6^{WT}$ + Doxo vs $mSTAT6^{WT}$ + Doxo + AZD2098, p = 0.9994<br>Post: $mSTAT6^{D419N}$ + Doxo vs $mSTAT6^{D419N}$ + Doxo + AZD2098, p = 0.0006<br>$mSTAT6^{WT}$ + Doxo: Pre vs Post, p = 0.3082<br>$mSTAT6^{D419N}$ + Doxo: Pre vs Post, p = 0.0135<br>$mSTAT6^{WT}$ + Doxo + AZD2098: Pre vs Post, p = 0.4645<br>$mSTAT6^{D419N}$ + Doxo + AZD2098: Pre vs Post, p = 0.6449 |
| 5I | Gehan-Breslow-Wilcoxon test, Bonferroni corrected | $mSTAT6^{WT}$ + Doxo = 5<br>$mSTAT6^{D419N}$ + Doxo = 5<br>$mSTAT6^{WT}$ + Doxo + AZD2098 = 5<br>$mSTAT6^{D419N}$ + Doxo + AZD2098 = 5 | $mSTAT6^{WT}$ + Doxo vs $mSTAT6^{D419N}$ + Doxo, p = 0.0070<br>$mSTAT6^{WT}$ + Doxo + AZD2098 vs $mSTAT6^{D419N}$ + Doxo + AZD2098, p = 0.0248<br>$mSTAT6^{WT}$ + Doxo vs $mSTAT6^{D419N}$ + Doxo + AZD2098, p = 0.1446<br>$mSTAT6^{D419N}$ + Doxo vs $mSTAT6^{D419N}$ + Doxo + AZD2098, p = 0.0095 |
| **Figure 6** | | | |
| 6G | Unpaired t test, two-tailed | $STAT6^{WT}$ = 31<br>$STAT6^{D419N}$ =6 | phospho-STAT6: $STAT6^{WT}$ vs $STAT6^{D419N}$, p < 0.0001<br>CCL17: $STAT6^{WT}$ vs $STAT6^{D419N}$, p < 0.0001 |
| 6H | Pearson correlation | Number of pairs = 37<br>$STAT6^{WT}$ = 31<br>$STAT6^{D419N}$ =6 | r = 0.8019<br>p < 0.0001 |
| 6I | Unpaired t test, two-tailed | $STAT6^{WT}$ = 31<br>$STAT6^{D419N}$ =6 | CCR4: $STAT6^{WT}$ vs $STAT6^{D419N}$, p = 0.0003 |
| 6J | Pearson correlation | Number of pairs = 37<br>$STAT6^{WT}$ = 31<br>$STAT6^{D419N}$ =6 | r = 0.5969<br>p < 0.0001 |
| 6K | Unpaired t test, two-tailed | $STAT6^{WT}$ = 31<br>$STAT6^{D419N}$ =6 | CD4 and Tumour: $STAT6^{WT}$ vs $STAT6^{D419N}$, p = 0.369 |

| | | | |
|---|---|---|---|
| | | | CCR4+ CD4 and Tumour: $STAT6^{WT}$ vs $STAT6^{D419N}$, p = 0.767<br>CD4 and pSTAT6+ Tumour: $STAT6^{WT}$ vs $STAT6^{D419N}$, p = 0.021<br>CCR4+ CD4 and pSTAT6+ Tumour: $STAT6^{WT}$ vs<br>$STAT6^{D419N}$, p = 0.124 |
| **Figure S1** | | | |
| S1A | Unpaired t test, two-tailed | D8 $mSTAT6^{WT}$ = 3<br>D8 $mSTAT6^{D419N}$ = 4 | % of CD45: Day 8 $mSTAT6^{WT}$ vs Day 8 $mSTAT6^{D419N}$, p = 0.7900<br>Number of cells: Day 8 $mSTAT6^{WT}$ vs Day 8 $mSTAT6^{D419N}$, p = 0.2898 |
| S1B | Unpaired t test, two-tailed | D14 $mSTAT6^{WT}$ = 5<br>D14 $mSTAT6^{D419N}$ = 4 | % of CD45: Day 14 $mSTAT6^{WT}$ vs Day 14 $mSTAT6^{D419N}$, p = 0.0934<br>Number of cells: Day 14 $mSTAT6^{WT}$ vs Day 14 $mSTAT6^{D419N}$, p = 0.1598 |
| S1C | Unpaired t test, two-tailed | D8 $mSTAT6^{WT}$ = 3<br>D8 $mSTAT6^{D419N}$ = 4 | % of CD45: Day 8 $mSTAT6^{WT}$ vs Day 8 $mSTAT6^{D419N}$, p = 0.8780<br>Number of cells: Day 8 $mSTAT6^{WT}$ vs Day 8 $mSTAT6^{D419N}$, p = 0.4364 |
| S1D | Unpaired t test, two-tailed | D14 $mSTAT6^{WT}$ = 5<br>D14 $mSTAT6^{D419N}$ = 4 | % of CD45: Day 14 $mSTAT6^{WT}$ vs Day 14 $mSTAT6^{D419N}$, p = 0.1788<br>Number of cells: Day 14 $mSTAT6^{WT}$ vs Day 14 $mSTAT6^{D419N}$, p = 0.2048 |
| **Figure S2** | | | |
| S2A | Unpaired t test, two-tailed | D11 $mSTAT6^{WT}$ = 10<br>D11 $mSTAT6^{D419N}$ = 10 | Day 11 $mSTAT6^{WT}$ vs Day 11 $mSTAT6^{D419N}$, p = 0.3681 |
| S2B | Unpaired t test, two-tailed | D11 $mSTAT6^{WT}$ = 10<br>D11 $mSTAT6^{D419N}$ = 10 | Day 11 $mSTAT6^{WT}$ vs Day 11 $mSTAT6^{D419N}$, p = 0.7515 |
| S2C | Unpaired t test, two-tailed | D11 $mSTAT6^{WT}$ = 10<br>D11 $mSTAT6^{D419N}$ = 10 | Day 11 $mSTAT6^{WT}$ vs Day 11 $mSTAT6^{D419N}$, p = 0.3017 |
| S2D | Unpaired t test, two-tailed | D11 $mSTAT6^{WT}$ = 10<br>D11 $mSTAT6^{D419N}$ = 10 | Day 11 $mSTAT6^{WT}$ vs Day 11 $mSTAT6^{D419N}$, p = 0.8248 |
| S2E | Unpaired t test, two-tailed | D11 $mSTAT6^{WT}$ = 10<br>D11 $mSTAT6^{D419N}$ = 10 | Day 11 $mSTAT6^{WT}$ vs Day 11 $mSTAT6^{D419N}$, p = 0.7978 |
| S2F | Unpaired t test, two-tailed | D11 $mSTAT6^{WT}$ = 10<br>D11 $mSTAT6^{D419N}$ = 10 | Day 11 $mSTAT6^{WT}$ vs Day 11 $mSTAT6^{D419N}$, p = 0.6957 |
| **Figure S3** | | | |
| S3A | Unpaired t test, two-tailed | $mSTAT6^{WT}$ + 3x3 mg/kg Doxo = 4<br>$mSTAT6^{D419N}$ + 3x3 mg/kg Doxo = 4<br>$mSTAT6^{WT}$ + 10 mg/kg Doxo = 8<br>$mSTAT6^{D419N}$ + 10 mg/kg Doxo = 10 | 3x3 mg/kg Doxo: $mSTAT6^{WT}$ vs $mSTAT6^{D419N}$, p = 0.5393<br>10 mg/kg Doxo: $mSTAT6^{WT}$ vs $mSTAT6^{D419N}$, p = 0.8210 |
| S3B | Unpaired t test, two-tailed | $mSTAT6^{WT}$ + 3x3 mg/kg Doxo = 4<br>$mSTAT6^{D419N}$ + 3x3 mg/kg Doxo = 4 | 3x3 mg/kg Doxo: $mSTAT6^{WT}$ vs $mSTAT6^{D419N}$, p = 0.7304<br>10 mg/kg Doxo: $mSTAT6^{WT}$ vs $mSTAT6^{D419N}$, p = 0.1555 |

| | | | |
|---|---|---|---|
| | | mSTAT6$^{WT}$ + 10 mg/kg Doxo = 8<br>mSTAT6$^{D419N}$ + 10 mg/kg Doxo = 10 | |
| S3C | Unpaired t test, two-tailed | mSTAT6$^{WT}$ + 3x3 mg/kg Doxo = 4<br>mSTAT6$^{D419N}$ + 3x3 mg/kg Doxo = 4<br>mSTAT6$^{WT}$ + 10 mg/kg Doxo = 8<br>mSTAT6$^{D419N}$ + 10 mg/kg Doxo = 10 | 3x3 mg/kg Doxo: mSTAT6$^{WT}$ vs mSTAT6$^{D419N}$, $p = 0.7585$<br>10 mg/kg Doxo: mSTAT6$^{WT}$ vs mSTAT6$^{D419N}$, $p = 0.8938$ |
| S3D | Unpaired t test, two-tailed | mSTAT6$^{WT}$ + 3x3 mg/kg Doxo = 4<br>mSTAT6$^{D419N}$ + 3x3 mg/kg Doxo = 4<br>mSTAT6$^{WT}$ + 10 mg/kg Doxo = 8<br>mSTAT6$^{D419N}$ + 10 mg/kg Doxo = 10 | 3x3 mg/kg Doxo: mSTAT6$^{WT}$ vs mSTAT6$^{D419N}$, $p = 0.7290$<br>10 mg/kg Doxo: mSTAT6$^{WT}$ vs mSTAT6$^{D419N}$, $p = 0.5891$ |
